# FAM84B facilitates tau propagation via RYR3-mediated exocytosis in response to neuroinflammation

**DOI:** 10.64898/2025.12.28.696549

**Authors:** Gwangho Yoon, Hyeyeon Park, Jee-Yun Park, Min Kyoung Kam, Jung-Hyun Kim, Eun-Joo Kim, Sung-Hye Park, Hyunyoung Kim, Gail VW Johnson, Ji-Young Choi, Young Ho Koh, Chulman Jo

**Affiliations:** Division of Brain Disease Research, 187 Osongsaengmyeong2-ro, Osong-eup, Cheongju-si, Chungcheongbuk-do, 28159, Republic of Korea; Division of Allergy and Respiratory Disease Research, Department for Chronic Disease Convergence Research, Korea National Institute of Health, 187 Osongsaengmyeong2-ro, Osong-eup, Cheongju-si, Chungcheongbuk-do, 28159, Republic of Korea; Division of Intractable Disease Research, Department for Chronic Disease Convergence Research, Korea National Institute of Health, 202 Osongsaengmyeong2-ro, Osong-eup, Cheongju-si, Chungcheongbuk-do, 28160, Republic of Korea; Korea National Stem Cell Bank, Korea National Institute of Health, 202 Osongsaengmyeong2-ro, Osong-eup, Cheongju-si, Chungcheongbuk-do, 28160, Republic of Korea; Adult Stem Cell Research Center and Research Institute for Veterinary Science, College of Veterinary Medicine, Seoul National University, Seoul, 08826, Republic of Korea; Department of Pharmacy, Ajou University, College of Pharmacy, 206 Worldcup-ro, Yeongtong-gu, Suwon-si 16499, Gyeonggi-do, Republic of Korea; Department of Neurology, Pusan National University Hospital, Pusan National University School of Medicine and Biomedical research institute, Busan, Republic of Korea; Department of Pathology, Seoul National University College of Medicine, Seoul, 03080, Republic of Korea; Department of Anesthesiology and Perioperative Medicine, University of Rochester, Rochester, NY, USA

**Author notes:** Correspondence and requests for materials should be addressed to C.J. These two authors equally contributed to this work (Gwangho Yoon, Hyeyeon Park).

## Abstract

The increasing prevalence of dementia underscores the pressing need for effective therapeutic interventions. Despite extensive research into Aβ, the limited efficacy of existing treatment modalities necessitates a paradigm shift toward tau pathology including tau propagation to slow dementia progression. Here, we identified a novel protein, FAM84B (also known as LRATD2), involved in tau propagation. Elevated FAM84B levels were detected in the postmortem cortex, cerebrospinal fluid, and induced pluripotent stem cell-derived cortical neurons of patients with tauopathy. FAM84B facilitates tau secretion through exosome assembly and RYR3-mediated exocytosis, followed by exosomal uptake into neighboring cells via clathrin-mediated endocytosis, thereby driving tau propagation. We confirmed that neuroinflammation induced by heightened STAT3 activity and cytokine treatment upregulated FAM84B expression, stimulating tau propagation, as evidenced by reduced propagation upon RYR3 inhibition. These findings provide invaluable insights into potential avenues for targeting tau propagation to ameliorate disease progression in tauopathies.

## Introduction

The rising prevalence of dementia and persistent challenges in developing effective treatments underscore the urgent need for a deeper understanding of the underlying causes and therapeutic targets^1^. Despite the fact that there have been significant research efforts focused on Aβ as a therapeutic target in Alzheimer’s disease (AD), results of clinical intervention studies targeting this peptide have been disappointing^2, 3^. This reflects the complexity of the pathophysiology of dementia and highlights the necessity for a more nuanced understanding of disease mechanisms as well as the exploration of alternative treatments^4^.

Addressing tau pathology is crucial for advancing the field of dementia research. Aberrant modifications of the tau protein, including hyperphosphorylation and aggregation, have been observed in various neurodegenerative disorders, including corticobasal degeneration (CBD), progressive supranuclear palsy (PSP), primary age-related tauopathy (PART), frontotemporal dementia (FTD), and AD, and are likely to contribute to cognitive decline^5^. Although interventions targeting tau pathology are promising therapeutic advancements, effective treatments currently remain elusive^6^. Strategies that focus on inhibiting tau aggregation, enhancing tau clearance mechanisms, or modulating tau-related cellular pathways provide potential avenues for slowing or halting disease progression, offering hope for developing effective treatments to alleviate the burden of dementia^7–10^.

Tau propagation has emerged as a critical focus area in recent studies shedding light on the complex nature of neurodegenerative conditions^11^, revealing diverse mechanisms facilitating the transmission of abnormal tau proteins between cells such as synaptic activity, exosome formation, transmission via nanotube formation, and tau-specific receptor-mediated endocytosis^12–14^. The pathophysiological complexities pose significant hurdles to therapeutic advancements in the field^15^. Although targeting tau aggregation within neurons and preventing its spread to neighboring cells are key strategies, identifying the specific genes or proteins responsible for tau propagation remains a challenge^16^.

In the present study, we identified FAM84B, also known as LRATD2, as a pivotal player in the pathological propagation of tau and linked it to neuroinflammation. Our findings suggest that FAM84B is a promising therapeutic target for attenuating disease progression in dementia. As well as representing a significant advancement in our understanding of dementia progression, these findings offer valuable insights to guide the development of innovative treatment strategies for tauopathies such as AD.

## Results

### FAM84B levels increase in association with tauopathy

HEK 293 cell lines stably expressing wild-type tau (2N4R) were prepared as a cell model of tauopathy in the presence of G418 antibiotics. Among these cell lines, we identified one with high intracellular accumulation of tau protein; to determine why these cells had different physiological properties, we analyzed their gene expression profiles. Microarray analysis of RNA isolated from the cells identified several genes with a differential expression (DEG) compared to control cells with relatively low tau expression levels (Supplementary Fig. 1). Notably, FAM84B (also known as LRATD2) expression decreased by approximately 18-fold in high tau-expressing cells compared to that in low tau-expressing cells. The involvement of FAM84B in tauopathy, including AD, has not been previously reported.

We further investigated FAM84B expression in various postmortem cortical brain tissues and cerebrospinal fluid (CSF) samples from patients with neurodegenerative diseases, including vascular dementia (VD), CBD, PSP, PART, mild cognitive impairment (MCI), and AD. A significant increase in FAM84B expression was observed in the postmortem cerebral cortex of patients with CBD, PSP, PART, and AD, but not in those with VD, compared to controls (Fig. 1A and Supplementary Fig. 2A, B). Additionally, elevated FAM84B levels were detected in the CSF of patients with MCI and AD compared with controls (Fig. 1B). These findings indicated the existence of a positive correlation between FAM84B expression and tauopathy-associated neurodegeneration.

**Figure 1.**
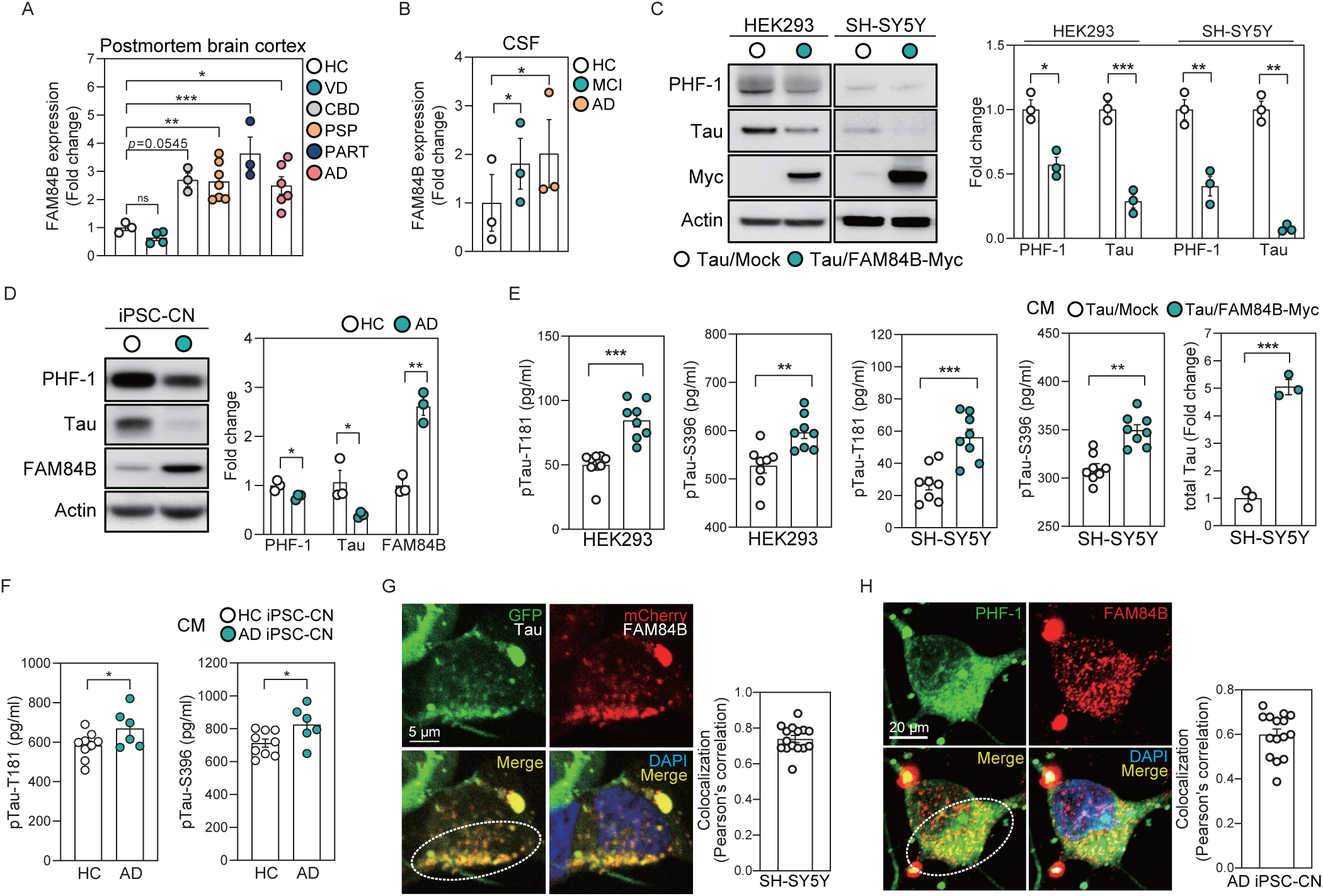
FAM84B mediates the secretion of phosphorylated and total tau proteins. **(A)** FAM84B levels in the cerebral cortex of HC (n=3) and patients with VD (n=4), CBD (n=3), PSP (n=7), PART (n=3), and AD (n=6). **(B)** FAM84B levels in the CSF of HC and patients with MCI and AD (n=3 each). **(C)** Levels of PHF-1, total tau, and FAM84B-Myc in HEK293 and SH-SY5Y cells expressing tau alone or tau with FAM84B-Myc (n=3 each). **(D)** Levels of PHF-1, total tau, and FAM84B in iPSC-derived CN from HC and patients with AD (n=3 each). **(E)** Levels of phosphorylated tau (T181 and S396) and total tau in the CM from HEK293 and SH-SY5Y cells expressing tau alone or tau with FAM84B-Myc (n=8 each). **(F)** Levels of phosphorylated tau (T181 and S396) in the CM from iPSC-derived CN from HC (n=9) and patients with AD (n=6). **(G)** Colocalization (dashed circle) of tau and FAM84B in SH-SY5Y cells expressing GFP-tau and mCherry-FAM84B (n=15). **(H)** Colocalization (dashed circle) of tau and FAM84B in iPSC-derived CN from patients with AD (n=15). Results are presented as mean ± SEM. Co-localization analysis: (G, H) Pearson’s correlation coefficient. Statistical analyses: (A, B) ordinary two-way ANOVA; (C–F) unpaired two-tailed *t*-test with Welch’s correction; *p < 0.05, **p < 0.01, and ***p < 0.001. Abbreviations: HC, healthy controls; VD, vascular dementia; CBD, corticobasal degeneration; PSP, progressive supranuclear palsy; PART, primary age-related tauopathy; AD, Alzheimer’s disease; CSF, cerebrospinal fluid; MCI, mild cognitive impairment; CN, cortical neurons; CM, conditioned media; PHF-1, phosphorylated tau at residues S396/S404

### FAM84B prompts tau secretion

To investigate the effect of increased FAM84B expression on tauopathy, HEK293 cells were transfected with varying amounts of tau and FAM84B. Increased FAM84B levels significantly reduced intracellular phosphorylated (PHF-1, S396/S404) and total tau protein levels in tau-expressing cells, irrespective of the quantity of tau introduced (Fig. 1C and Supplementary Fig. 3A). Additional experiments using mouse neuroblastoma cells (Neuro-2A) and human neuroblastoma cells (SH-SY5Y), as well as their differentiated forms, also showed that FAM84B overexpression led to a significant decrease in intracellular phosphorylated and total tau protein levels in all tested cell models (Fig. 1C and Supplementary Fig. 3B).

Next, we used cortical neurons derived from induced pluripotent stem cells (iPSCs) obtained from healthy individuals and patients with AD to confirm the role of FAM84B in neurons. Elevated levels of FAM84B were observed alongside decreased intracellular levels of phosphorylated and total tau proteins in iPSC-derived cortical neurons from patients with AD compared to those from healthy individuals (Fig. 1D). These findings suggest that FAM84B is involved in the regulation of tau proteins in all examined cell models, including iPSC-derived neurons.

Further exploration of FAM84B’s influence on intracellular tau protein levels revealed that the reduction in tau protein levels mediated by FAM84B was not owing to transcriptional or degradative regulation. FAM84B overexpression did not affect MAPT mRNA levels in tau-expressing cells (Supplementary Fig. 4A). Elevated FAM84B levels consistently decreased total tau protein levels in a doxycycline-inducible tau (DIT) overexpression system in mouse cortical neurons (Supplementary Fig. 4B). Additionally, increased levels of FAM84B did not affect the levels of proteins associated with autophagy (p62 and LC3) and lysosomal degradation (HSP70, LAMP-1, and LAMP-2), a well-known pathway for tau degradation, in tau-expressing cells (Supplementary Fig. 4C, D).

Notably, we observed that FAM84B promotes the secretion of phosphorylated and total tau proteins from tau-expressing cells (Fig. 1E). Tau protein was secreted along with FAM84B (Supplementary Fig. 5A). An increase in phosphorylated tau protein secretion was also observed in iPSC-derived cortical neurons from patients with AD compared to that in healthy individuals (Fig. 1F). Additionally, FAM84B reduced the intracellular levels of phosphorylated and total P301L mutant tau while enhancing its release into the extracellular space (Supplementary Fig. 5B–D). We confirmed that FAM84B induces comparable levels of tau protein secretion in wild-type and P301L mutant tau (Supplementary Fig. 5E), indicating that FAM84B does not distinguish between tau species during secretion. These findings suggest that the reduction in intracellular tau protein levels associated with FAM84B occurs through extracellular secretion, emphasizing FAM84B’s significant involvement in the tau protein secretion pathway.

To investigate the effects of FAM84B deletion on tau protein secretion, FAM84B knockout HEK293 cells were generated (Supplementary Fig. 6A). Overexpression of wild-type tau or P301L mutant tau in FAM84B knockout cells led to a decrease in the extracellular levels of total and phosphorylated tau proteins across all tau variants (Supplementary Fig. 6B, C). Conversely, FAM84B knockout cells exhibited intracellular accumulation of all tau variants (Supplementary Fig. 6D). Taken together, these results strongly suggest a crucial role for FAM84B in tau secretion that is independent of tau phosphorylation and species.

### FAM84B mediates exosomal tau propagation

To investigate the potential colocalization of FAM84B and tau proteins during secretion, we used various tauopathy cell models with plasmids tagged with fluorescent genes or via immunocytochemistry techniques. In cells expressing tau-GFP and FAM84B-mCherry, we observed their colocalization as intracellular and extracellular granules (Fig. 1G and Supplementary Fig. 7A, B, G). Immunocytochemistry further confirmed the significant co-localization between FAM84B and phosphorylated and total tau proteins across multiple cell models, including iPSC-derived cortical neurons from patients with AD (Fig. 1H and Supplementary Fig. 7C–G). These findings suggest that tau secretion in association with FAM84B likely occurs via intracellular vesicle formation or protein-protein interactions.

Immunoprecipitation assays were performed using lysates and culture media from cells overexpressing tau and FAM84B, to explore the presence of interactions. No direct interaction between FAM84B and tau was detected in cell lysates prepared in the presence of the detergent NP-40 (Supplementary Fig. 8A). However, high molecular weight tau coprecipitated with FAM84B in culture media under detergent-free conditions (Supplementary Fig. 8B). This high molecular weight tau consisted entirely of monomeric tau following the boiling of the precipitates in the presence of a reducing agent (Supplementary Fig. 8C). These results suggest that FAM84B and tau coexist within the lipid compartment in the culture medium disrupted by detergents. Upon lipid component disruption using NP-40, no FAM84B-tau precipitates were observed (Supplementary Fig. 8D), indicating their coexistence within extracellular vesicles, such as exosomes.

To confirm the involvement of exosomes in the secretion of FAM84B and tau, we isolated exosomes from the culture media of cells overexpressing FAM84B and tau. The presence of the exosomal marker Alix indicated that FAM84B, phosphorylated tau, and total tau proteins were encapsulated in exosomes (Fig. 2A, B). Membrane protein isolation revealed that FAM84B is a component of the exosome membrane along with caveolin-1 (Fig. 2C), suggesting that FAM84B facilitates tau secretion by assembling exosome membranes that encapsulate tau proteins.

**Figure 2.**
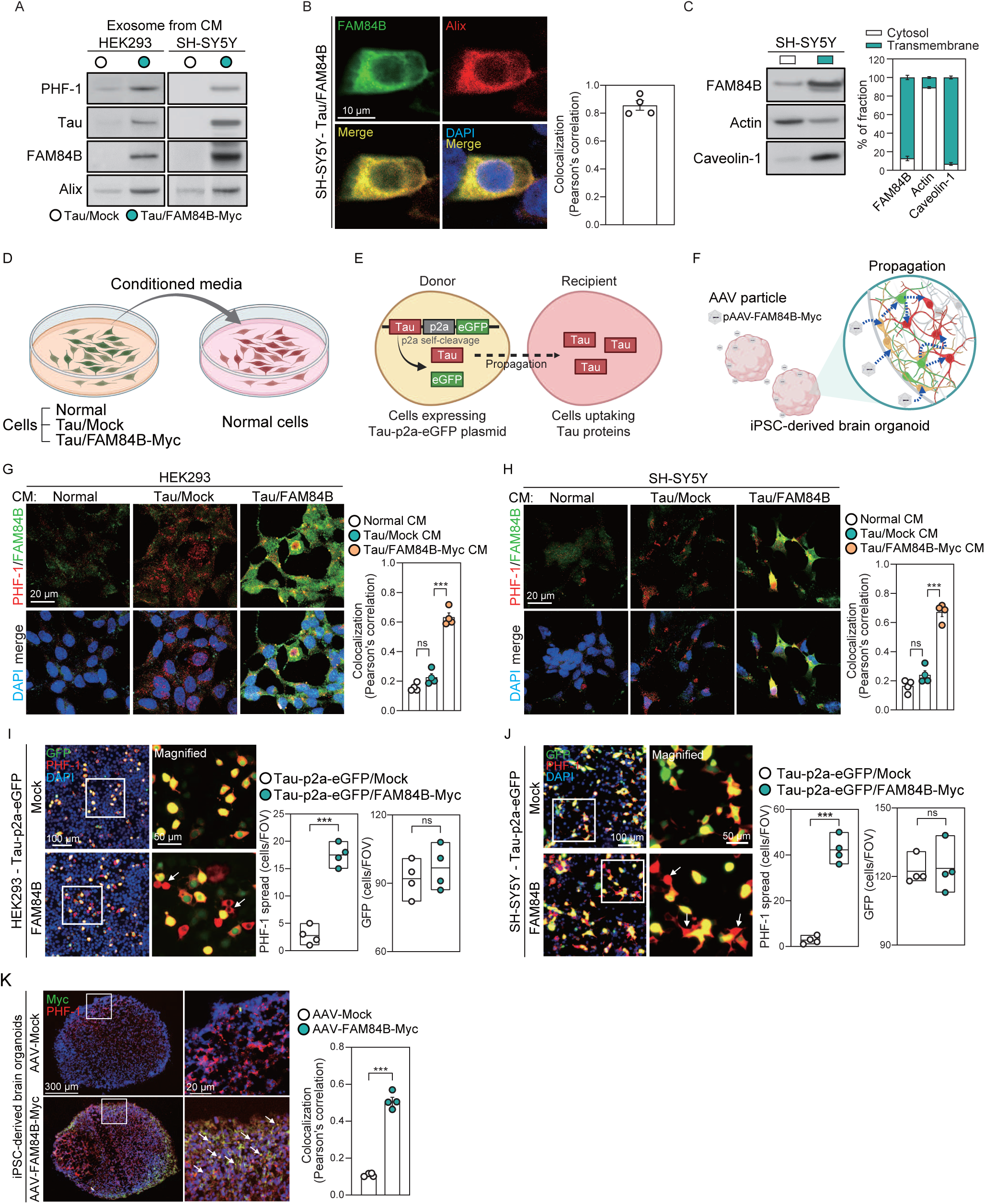
FAM84B promotes tau propagation via exosomal formation. **(A)** Immunoblots showing levels of PHF-1, total tau, FAM84B, and Alix in exosomes from the CM of HEK293 and SH-SY5Y cells expressing tau alone or tau with FAM84B-Myc (n=3 each). **(B)** Colocalization of FAM84B and Alix in SH-SY5Y cells expressing tau with FAM84B (n=4). **(C)** Relative levels of FAM84B, actin, and caveolin-1 in cytosolic and transmembrane fractions of SH-SY5Y cells expressing tau with FAM84B (n=3). **(D)** Illustration describing the administration of CM from normal cells, cells expressing tau alone, and cells expressing tau with FAM84B-Myc into normal cells, supporting the results shown in (G) and (H). **(E)** Illustration depicting tau transmission from Tau-p2a-eGFP expressing cells (donor) to neighboring cells (recipient), supporting the results shown in (I) and (J). **(F)** Illustration showing tau propagation in iPSC-derived brain organoids from HC infected with AAV particles containing pAAV-FAM84B-Myc, pAAV-Tau-Myc-p2a-eGFP, or pAAV-FAM84B, supporting the results shown in (K). **(G)** Uptake and colocalization of PHF-1 and FAM84B in HEK293 cells exposed to CM from normal cells, cells expressing tau alone, and cells expressing tau with FAM84B-Myc (n=4 each). **(H)** Uptake and colocalization of PHF-1 and FAM84B in SH-SY5Y cells exposed to CM from normal cells, cells expressing tau alone, and cells expressing tau with FAM84B-Myc (n=4 each). **(I)** PHF-1 spread from Tau-p2a-eGFP expressing HEK293 cells to neighboring cells in response to FAM84B expression (n=4 each). Arrows indicate recipient cells (red) to which tau has been transmitted. Solid boxes in the images indicate magnified areas. **(J)** PHF-1 spread from Tau-p2a-eGFP expressing SH-SY5Y cells to neighboring cells in response to FAM84B expression (n=4 each). Arrows indicate recipient cells (red) to which tau has been transmitted. Solid boxes in the images indicate magnified areas. **(K)** Colocalization (arrow) of PHF-1 and FAM84B (Myc) in iPSC-derived brain organoids from HC infected with AAV containing control vector or FAM84B-Myc (n=4 each). Solid boxes in the images indicate magnified areas. Results are presented as mean ± SEM. Co-localization analysis: (B, G, H, K) Pearson’s correlation coefficient. Statistical analyses: (G–K) Unpaired two-tailed *t*-test with Welch’s correction; ns: not significant; ***p < 0.001. Illustration (D-F) was created with BioRender.com (Agreement number: XO2789HRIK, RF2789HV9J, VS2789HBYY) Abbreviations: PHF-1, phosphorylated tau at residues S396/S404; CM, conditioned media; HC, healthy control; AAV, adeno-associated virus; FOV, field of view.

To further elucidate the role of FAM84B in tau propagation, we employed three approaches to investigate FAM84B-mediated tau transmission to neighboring cells (Fig. 2D–F). First, we exposed normal cells to culture medium containing tau secreted by FAM84B cells to determine whether tau uptake occurred (Fig. 2D). Our findings showed a significantly higher uptake of phosphorylated and total tau in HEK293 and SH-SY5Y cells exposed to secretions from cells co-expressing FAM84B and tau than in those exposed to secretions from cells expressing tau alone (Fig. 2G, H, and Supplementary Fig. 9A–C). Phosphorylated and total tau exhibited high co-localization with FAM84B in cells treated with co-expressed secretions compared to tau-only secretions.

Next, we investigated tau transmission to neighboring cells when FAM84B was overexpressed in cells transfected with a tau propagation plasmid (Tau-p2a-eGFP) (Fig. 2E). A significant increase in the number of cells exhibiting tau uptake was observed when FAM84B was overexpressed in Tau-p2a-eGFP-transfected cells compared to cells lacking FAM84B expression (Fig. 2I, J).

Finally, we evaluated tau propagation within brain organoids infected with recombinant adeno-associated virus (AAV) containing the FAM84B gene (Fig. 2F). The outer region of the brain organoids exposed to AAV particles containing FAM84B exhibited significant FAM84B expression and prominent colocalization between FAM84B and phosphorylated tau compared to the control (Fig. 2K). Collectively, these analyses indicated that FAM84B plays a pivotal role in tau propagation through exosome-mediated secretion.

### FAM84B induces ryanodine receptor 3 (RYR3)/clathrin-mediated tau propagation

To investigate the mechanism underlying FAM84B-mediated exosome secretion, we performed mRNA-seq analysis using RNA extracted from cells with FAM84B deletion, FAM84B overexpression, or simultaneous overexpression of FAM84B and tau (Fig. 3A–C). This identified common genes among the altered genes in each cell type, revealing changes related to intracellular plasma membrane constituents consistent with alterations in FAM84B expression (Fig. 3D, E). These findings suggest that FAM84B regulates the genes involved in intracellular membrane dynamics, shedding light on its role in exosome secretion.

**Figure 3.**
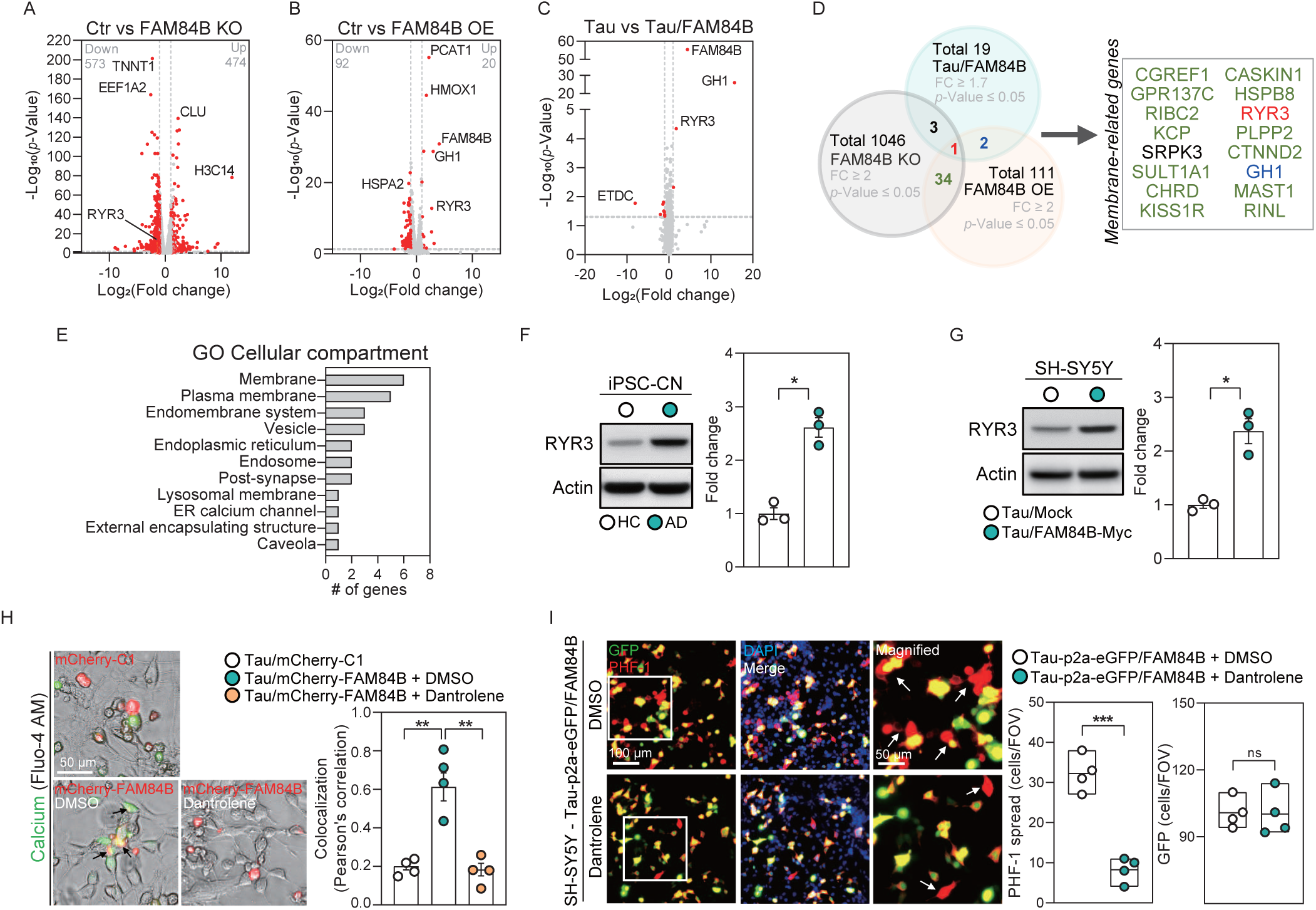
FAM84B induces RYR3-mediated tau propagation. **(A)** Differentially expressed genes in HEK293 cells in response to FAM84B KO (n=3 each). Genes with p < 0.05 and >2-fold expression change compared to control are marked with red dots. **(B)** Differentially expressed genes in HEK293 cells in response to FAM84B OE (n=3 each). Genes with p < 0.05 and >2-fold expression change compared to control are marked with red dots. **(C)** Differentially expressed genes in tau-expressing SH-SY5Y cells in response to FAM84B OE (n=3 each). Genes with p < 0.05 and >1.7-fold expression change compared to cells expressing tau alone are marked with red dots. **(D)** Venn diagram showing the number of shared genes between RNA-seq datasets in (A–C), with overlapping genes represented in distinct colors (red, green, black, blue). Only the names of membrane-related genes among the shared genes are noted. **(E)** GO analysis showing cellular compartments of genes identified in (D). The GO terms based on the number of genes are shown. **(F)** Levels of RYR3 and actin in iPSC-derived CN from HC and patients with AD (n=3 each). **(G)** Levels of RYR3 and actin in SH-SY5Y cells expressing tau alone or tau with FAM84B-Myc (n=3 each). **(H)** Colocalization (arrow) of FAM84B and calcium in SH-SY5Y cells expressing tau with mCherry control (C1), and in cells expressing tau with mCherry-FAM84B treated with either DMSO or dantrolene (n=4 each). **(I)** PHF-1 spread from SH-SY5Y cells expressing Tau-p2a-eGFP with FAM84B to neighboring cells in response to treatment with either DMSO or dantrolene (n=4 each). Arrows indicate recipient cells (red) to which tau has been transmitted. Solid boxes in the images indicate magnified areas. Results are presented as mean ± SEM. Co-localization analysis. (H) Pearson correlation coefficient. Statistical analyses: (A-C, F-I) unpaired two-tailed *t*-test with Welch’s correction; ns: not significant, *p < 0.05, **p < 0.01, ***p < 0.001. Abbreviations: KO, knockout; OE, overexpression; GO, gene ontology; CN, cortical neuron; HC, healthy control; AD, Alzheimer’s disease; PHF-1, phosphorylated tau at residues S396/S404; FOV, field of view.

Among the genes with altered expression patterns, RYR3 consistently appeared in all three mRNA-Seq datasets. This observation was corroborated through analysis of mRNA-seq databases, which demonstrated a positive correlation between FAM84B and RYR3 expression levels (r = 0.97). In cortical neurons derived from iPSCs of patients with AD, we observed a significant increase in RYR3 levels compared to that in healthy controls (Fig. 3F). In a similar manner, there was a notable increase in RYR3 levels among SH-SY5Y cells overexpressing tau and FAM84B relative to that in cells expressing tau alone (Fig. 3G).

Previous studies have highlighted RYR3’s function as a calcium channel within the endoplasmic reticulum that regulates exocytosis by modulating cytosolic calcium levels^17, 18^. Staining for cytosolic calcium using Fluo-4 AM revealed a substantial increase in FAM84B–cytosolic calcium colocalization in cells co-expressing tau and mCherry-FAM84B compared to those co-expressing tau and mCherry-control (Fig. 3H and Supplementary Fig. 10A). This colocalization diminished markedly following treatment with the specific RYR3 inhibitor dantrolene, indicating that RYR3 upregulation by FAM84B is linked to elevated cytosolic calcium levels.

To investigate whether FAM84B-mediated tau propagation is RYR3-dependent, we treated cells coexpressing Tau-p2a-eGFP and FAM84B with dantrolene. A significant decrease in FAM84B-mediated tau propagation was observed following dantrolene treatment (Fig. 3I and Supplementary Fig. 10B). These findings suggest that the increased RYR3 expression induced by FAM84B triggers FAM84B-mediated tau exocytosis and propagation via the elevation of cytosolic calcium levels.

To explore the mechanism underlying the internalization of exosomes containing FAM84B and tau proteins by neighboring cells, the levels of endocytosis-associated proteins such as clathrin and caveolin and endosomal markers in cells co-expressing FAM84B and tau were compared to the levels in cells expressing tau alone. Elevated FAM84B expression significantly upregulated clathrin heavy chain (CHC) levels and downregulated caveolin-1 levels in tau-expressing cells compared to cells expressing tau alone (Fig. 4A). However, there were no discernible changes observed in early or late endosomal marker proteins (EEA1, RAB5, RAB7, RAB9, and RAB11) compared to controls. Increased levels of CHC were also evident in the postmortem cerebral cortex of patients with tauopathy, including those with PSP and AD, and in iPSC-derived cortical neurons of patients with AD compared to their respective normal controls (Fig. 4B, C), indicating a correlation between FAM84B and increased CHC levels in tauopathy-afflicted brains and neurons.

**Figure 4.**
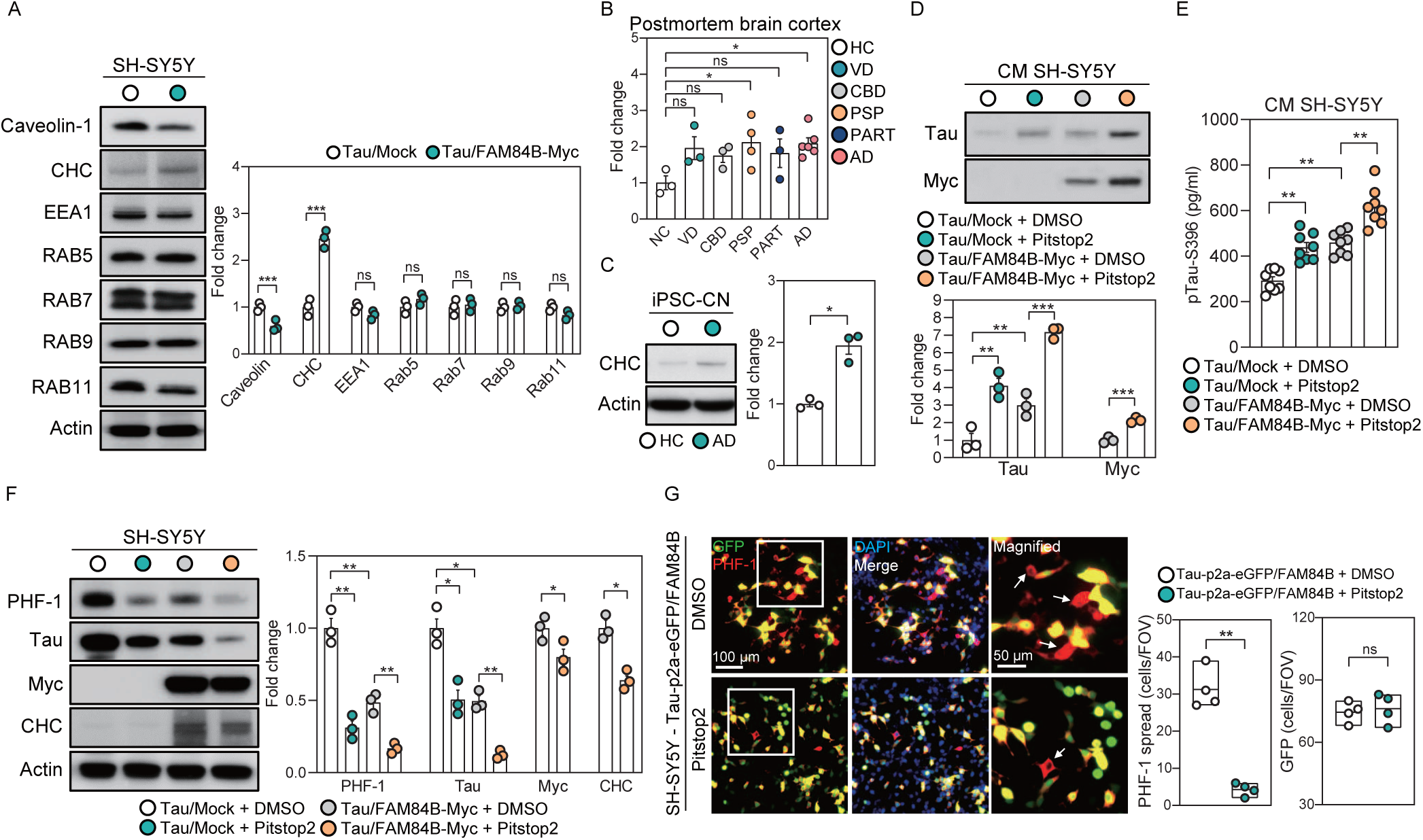
FAM84B induces clathrin-mediated tau propagation. **(A)** Levels of caveolin-1, CHC, EEA1, RAB5, RAB7, RAB9, RAB11, and actin in SH-SY5Y cells expressing tau alone or tau with FAM84B-Myc (n=3 each). **(B)** CHC levels in the cerebral cortex of HC (n=3) and patients with VD (n=3), CBD (n=3), PSP (n=4), PART (n=3), and AD (n=6). **(C)** Levels of CHC and actin in iPSC-derived CN from HC and patients with AD (n=3 each). **(D)** Changes in the levels of tau and FAM84B (Myc) in the CM from SH-SY5Y cells expressing tau alone or tau with FAM84B-Myc after treatment with either DMSO or Pitstop2 (n=3 each). **(E)** Changes in the levels of phosphorylated tau (S396) in the CM from SH-SY5Y cells expressing tau alone or tau with FAM84B-Myc after treatment with either DMSO or Pitstop2 (n=8 each). **(F)** Changes in the levels of PHF-1, total tau, FAM84B (Myc), CHC, and actin in SH-SY5Y cells expressing tau alone or tau with FAM84B-Myc after treatment with either DMSO or Pitstop2 (n=8 each). **(G)** PHF-1 spread from SH-SY5Y cells expressing Tau-p2a-eGFP with FAM84B to neighboring cells in response to treatment with either DMSO or Pitstop2 (n=4 each). Arrows indicate recipient cells (red) to which tau has been transmitted. Solid boxes in the images indicate magnified areas. Results are presented as mean ± SEM. Statistical analyses: (B) ordinary two-way ANOVA; (A, C-G) unpaired two-tailed *t*-test with Welch’s correction; ns: not significant, *p < 0.05, **p < 0.01, ***p < 0.001. Abbreviations: CHC, clathrin heavy chain; HC, healthy control; VD, vascular dementia; CBD, corticobasal degeneration; PSP, progressive supranuclear palsy; PART, primary age-related tauopathy; AD, Alzheimer’s disease; CN, cortical neuron; CM, conditioned medium; PHF-1, phosphorylated tau at S396/S404 residue; FOV, field of view.

To ascertain whether FAM84B-mediated tau uptake occurs via clathrin-mediated endocytosis, we used the clathrin-mediated endocytosis inhibitor pitstop2. Compared to untreated controls, pitstop2-treated cells either co-expressing tau and FAM84B or expressing tau alone exhibited elevated levels of extracellular phosphorylated tau and total tau (Fig. 4D, E), along with reduced levels of intracellular phosphorylated tau and total tau (Fig. 4F). Similarly, relative to untreated controls, pitstop2-treated cells coexpressing FAM84B and tau demonstrated increased extracellular FAM84B levels and decreased intracellular FAM84B levels (Fig. 4D, F). This indicates that exosomes containing FAM84B and tau were taken up by neighboring cells via clathrin-mediated endocytosis.

We validated the role of clathrin in FAM84B-mediated tau propagation in cells co-expressing Tau-p2a-eGFP and FAM84B. Following pitstop2 treatment, which inhibited clathrin-mediated endocytosis, a significant reduction in FAM84B-mediated tau propagation was observed compared to that in untreated controls (Fig. 4G). These findings indicate the pivotal role of clathrin-mediated endocytosis in facilitating FAM84B-mediated tau propagation.

### Neuroinflammation triggers FAM84B-mediated tau propagation

Intrigued by the observed upregulation of FAM84B in the postmortem cerebral cortex of tauopathy patients (Fig. 1A), we analyzed ChIP-seq data (ENCODE, ChEA) for transcription factors binding to the FAM84B promoter in order to identify the regulators of FAM84B expression. We identified transcription factors involved in inflammation (STAT3 and JUN), hormone signaling (ESR1), and chromatin remodeling (SUZ12, RNF2, ZNF217, and PHF8) as potential regulators (Fig. 5A). STAT3 binding was observed in association with the FAM84B enhancer, whereas ESR1 and JUN were associated with nearby regulatory elements (Fig. 5B). Increased STAT3 activity via Y705 phosphorylation was noted in the postmortem cerebral cortex of patients with tauopathy and iPSC-derived cortical neurons from patients with AD compared with controls (Fig. 5C, D and Supplementary Fig. 11A, B).

**Figure 5.**
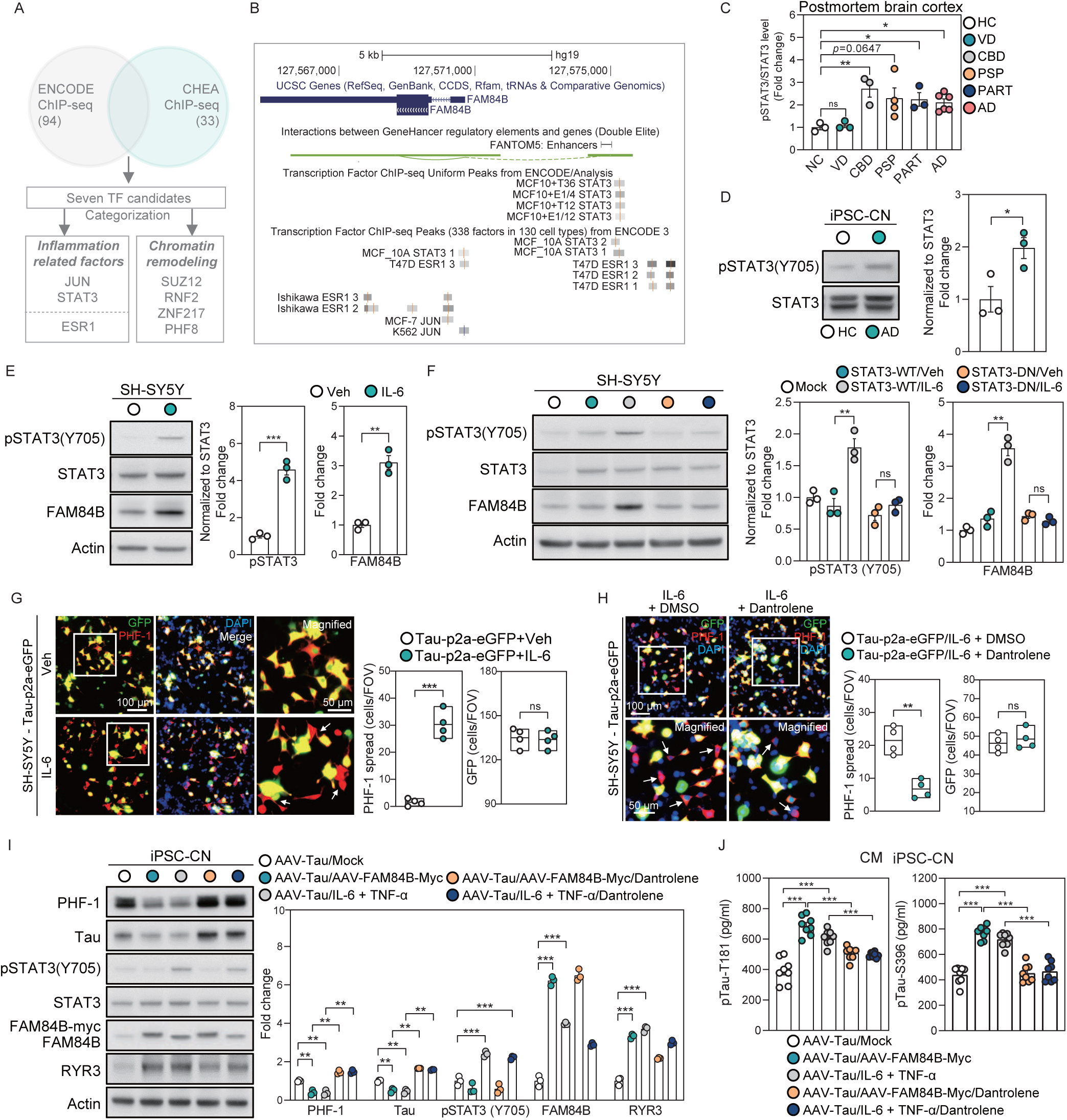
Neuroinflammation may trigger FAM84B-mediated tau propagation. **(A)** Venn diagram showing the seven shared transcriptional cofactors between the ENCODE ChIP-seq and CHEA ChIP-seq datasets. The genes are categorized into functional terms and noted. **(B)** Genomic information of the FAM84B gene obtained from the UCSC Genome Browser. Transcription cofactor clusters identified from the ENCODE ChIP-seq data, with those likely to bind to the regulatory elements of FAM84B, as identified by FANTOM5, highlighted. **(C)** Levels of phosphorylated STAT3 (Y705) in the cerebral cortex of HC (n=3) and patients with VD (n=3), CBD (n=3), PSP (n=4), PART (n=3), and AD (n=6). **(D)** Levels of phosphorylated STAT3 (Y705) and total STAT3 in iPSC-derived CN from HC and patients with AD (n=3 each). **(E)** Levels of phosphorylated STAT3 (Y705), total STAT3, FAM84B, and actin in SH-SY5Y cells treated with either control vehicle or IL-6 (n=3 each). **(F)** Changes in the levels of phosphorylated STAT3 (Y705), total STAT3, FAM84B, and actin in SH-SY5Y cells expressing STAT3-WT or STAT3-DN in response to treatment with either control vehicle or IL-6 (n=3 each). **(G)** PHF-1 spread from Tau-p2a-eGFP expressing SH-SY5Y cells to neighboring cells in response to treatment with either control vector or IL-6 (n=4 each). Arrows indicate recipient cells (red) to which tau has been transmitted. Solid boxes in the images indicate magnified areas. **(H)** PHF-1 spread from IL-6 treated SH-SY5Y cells expressing Tau-p2a-eGFP to neighboring cells in response to treatment with either DMSO or dantrolene (n=4 each). Arrows indicate recipient cells (red) to which tau has been transmitted. Solid boxes in the images indicate magnified areas. **(I)** Changes in the levels of PHF-1, total tau, phosphorylated STAT3 (Y705), total STAT3, FAM84B (FAM84B-Myc), RYR3, and actin in iPSC-derived CN from HC after infection with AAV and subsequent treatments (n=3 each). **(J)** Changes in the levels of phosphorylated tau (T181 and S396) in the CM of iPSC-derived CN from HC after infection with AAV and subsequent treatments (n=8 each). In (I–J), the experimental groups are: (1) CN infected with AAV containing tau, (2) CN infected with AAV containing tau and FAM84B-Myc, (3) CN infected with AAV containing tau and treated with IL-6 and TNF-α, (4) CN infected with AAV containing tau and FAM84B-Myc and treated with dantrolene, and (5) CN infected with AAV containing tau, treated with IL-6 and TNF-α, followed by treatment with dantrolene. Results are presented as mean ± SEM. Statistical analyses: (C) ordinary two-way ANOVA; (D-J) unpaired two-tailed *t*-test with Welch’s correction; ns: not significant, *p < 0.05, **p < 0.01, and ***p < 0.001. Abbreviations: HC, healthy control; VD, vascular dementia; CBD, corticobasal degeneration; PSP, progressive supranuclear palsy; PART, primary age-related tauopathy; AD, Alzheimer’s disease; CN, cortical neuron; WT, wild type; DN, dominant negative; PHF-1, phosphorylated tau at S396/S404 residue; FOV, field of view; AAV, adeno-associated virus; CM, conditioned media.

Proinflammatory cytokines, and notably interleukin (IL)-6, reliably phosphorylate STAT3 at Y705^19^. Treatment of SH-SY5Y cells with IL-6 increased levels of STAT3 phosphorylation at Y705 and FAM84B compared to those in untreated controls (Fig. 5E). This effect was consistent in IL-6-treated cells expressing wild-type STAT3 but absent in cells with a dominant-negative STAT3 mutation, indicating that FAM84B expression is regulated by IL-6-mediated STAT3 activation (Fig. 5F). Tumor necrosis factor (TNF)-α was also associated with increased FAM84B levels in SH-SY5Y cells (Supplementary Fig. 11C). Levels of tau propagation were elevated in cells expressing Tau-p2a-eGFP and treated with IL-6 or TNF-α compared to that in controls (Fig. 5G and Supplementary Fig. 11D). These findings suggest that neuroinflammation regulates FAM84B expression and promotes tau propagation.

We then validated the interplay between neuroinflammation, FAM84B, RYR3, and tau propagation in SH-SY5Y cells and iPSC-derived cortical neurons. The treatment of SH-SY5Y cells expressing Tau-p2a-eGFP with IL-6, followed by treatment with the RYR3 inhibitor dantrolene, significantly suppressed the increased tau propagation observed in untreated controls (Fig. 5H). These results suggested a correlation between neuroinflammation and RYR3 calcium channels.

Further validation was performed by infecting iPSC-derived cortical neurons with AAV particles encoding either tau or FAM84B, or alternatively treating the cortical neurons with a combination of IL-6 and TNF-α followed by dantrolene. RYR3 levels increased significantly in iPSC-derived cortical neurons expressing FAM84B via AAV infection and in those co-treated with IL-6 and TNF-α (Fig. 5I). A reduction in intracellular phosphorylated and total tau levels was consistently observed in cells co-expressing tau and FAM84B, as well as in those treated with IL-6 and TNF-α (Fig. 5I); furthermore, there was a simultaneous increase in extracellular phosphorylated tau levels in these cells compared with their respective controls (Fig. 5J).

In conclusion, the above findings indicate that neuroinflammatory responses elevate FAM84B levels that subsequently influence signaling pathways involving the RYR3 calcium channel, thereby contributing to the propagation of tau (Supplementary Fig. 12).

## Discussion

In AD, cognitive decline correlates more directly with the accumulation of tau than with amyloid plaque burden^20, 21^. The conceptual framework of tau pathology spreading through the brain in a prion-like manner has been supported by Braak and Braak^22^, wherein tau pathology is observed in a consistent spatiotemporal pattern that is closely correlated with clinical disease stages. Accumulating evidence indicates that neuroinflammation contributes to neurodegeneration^23^ and significantly accelerates the aggregation and phosphorylation of tau^24,25^. Furthermore, reactive microglia are sufficient to drive tau pathology and their presence is correlated with the spread of tau pathology throughout the brain^26^. However, the involvement and role of neuroinflammation in the transneuronal propagation of tau remains to be elucidated. Here, we show the complex interactions between neuroinflammation and tau transmission, revealing a critical role in tau propagation for FAM84B, which is a protein traditionally linked to cancer^27^, and RYR3. The valuable insights offered by these results could inform potential therapeutic strategies targeting the propagation of tau observed in tauopathies such as AD.

In the current study, we found that FAM84B significantly enhanced the transmission of wild-type and mutant tau (P301L) independent of their phosphorylation status (Fig. 1 and Supplementary Fig. 5). To elucidate the mechanisms underlying this effect, the specificity of FAM84B was investigated. Our results indicate that FAM84B promotes encapsulation rather than directly binding to tau (Fig. 2 and Supplementary Fig. 8). This encapsulation process did not selectively target key phosphorylation sites (T181, T231, S262, S396, and S404; data not shown), suggesting that FAM84B facilitates tau encapsulation in the most general sense. The ability of FAM84B to promote the encapsulation of tau into exosomes in a broad manner highlights its potential importance as a fundamental regulator of the pathogenesis of tauopathy-related diseases such as AD. Our results also indicate that FAM84B reduces intracellular tau levels through the promotion of tau secretion via exosome formation rather than degradation (Fig. 2 and Supplementary Fig. 4). This pathophysiological mechanism contributes significantly to the spread of tauopathy, highlighting the pathogenic role of extracellular tau, which is facilitated by FAM84B^28^. The formation of exosomes containing FAM84B and tau may be influenced by the functional and structural characteristics of FAM84B, suggesting that it has a role in vesicular trafficking and exosome biogenesis^29^. In the current body of work, FAM84B did not promote GFP or amyloid-beta secretion but did facilitate the secretion of alpha-synuclein (data not shown), suggesting the specificity of FAM84B for intrinsically disordered proteins prone to intracellular aggregation. These findings suggest that FAM84B influences disease progression through mechanisms that extend beyond tau propagation. Further research is required to identify other proteins secreted by FAM84B and to elucidate the mechanisms through which FAM84B encapsulates intrinsically disordered proteins.

In addition to exosome formation, FAM84B mediates tau propagation through clathrin-mediated endocytosis (CME) (Fig. 4), with mechanisms that vary across different diseases^30^. In AD, CME facilitated by receptors such as LRP1 predominates. LRP1 knockdown has been shown to almost eliminate tau uptake via CME in neuroglioma cells and reduce tau spread in mouse models^14^, corroborating the critical role of CME in tau propagation. However, different pathways such as microtubule-associated transport may operate in PSP and CBD, as both diseases are characterized by 4-repeat tauopathies with a strong microtubule affinity^31, 32^. A distinct increase in FAM84B expression was observed in specific tauopathies, namely CBD, PSP, PART, and AD, but not in VD, where FAM84B levels remained stable (Fig. 1 and Supplementary Fig. 2). This differential expression suggests a tauopathy-specific role for FAM84B, which is supported by recent evidence indicating elevated FAM84B levels in patients with AD^33^. Thus, our data suggest that FAM84B may play a central role in tau propagation through CME in the pathogenesis of various tauopathies including PSP and CBD; however, further research is needed to elucidate the specific roles of FAM84B in different tauopathies.

Our study has some limitations. We have concluded that elevated FAM84B levels exacerbate tau pathology in tauopathies by promoting tau propagation. Generally, intracellular aggregation of pathological tau is observed as a common feature in tauopathies which are distinct from one another with respect to predominant tau isoforms (3R or 4R) that accumulate, cell types that have tau inclusions, and vulnerable brain regions^32^. Although observing elevated FAM84B levels in most brain tissues of patients with tauopathy (Supplementary Fig. 2), in our cell models we found increased FAM84B expression was associated with reduced intracellular tau levels owing to enhanced tau secretion (Fig. 1). In this regards, elevated FAM84B levels may not fully explain the accumulation of pathological tau seen in various cell types in the brain tissues of patients with tauopathy^34^. Instead, we speculate that FAM84B may support the redistribution of tau from intracellular to extracellular regions or to other cells, such as neurons and glia^32^. Given that the diversity of tau pathology in tauopathies including CBD, PSP and AD, the aggregation of pathological tau in glial cells or neurons could be caused by other concurrent, pathological events such as the impairment of autophagy^35^, the posttranslational modification of tau like acetylation^36^ or the mutation of tau gene^37^, independent of FAM84B-mediated propagation.

Mounting evidence suggests that the pathogenesis of AD is not restricted to the neuronal compartment, but instead includes strong interactions with immunological mechanisms in the brain. Misfolded aggregates of Aβ are thought to bind to microglia and astroglia, triggering microglial activation and leading to the release of inflammatory cytokines such as IL-6 and TNF-α that contribute to disease progression as well as severity^38, 39^. As shown in Fig. 5C, STAT3 was activated in the brain tissues of patients with tauopathy alongside increased FAM84B expression (Fig. 1A). In line with these results, IL-6 and TNF-α can induce FAM84B expression (Fig. 5G and Supplementary Fig. 11D). Given that FAM84B is a crucial mediator of tau propagation, these results strongly suggest that neuroinflammation induces FAM84B expression and stimulates tau propagation in the brain, leading to increased tau pathology. Thus, our results suggest that modulation or targeting of immune mechanisms could lead to future therapeutic or preventive strategies for tauopathies, including AD. In conclusion, by demonstrating the important role FAM84B plays in tau propagation and intercellular communication during neuroinflammation, our study significantly enhances the understanding of tauopathies. Integration and application of these insights could enhance the development of targeted therapies to effectively address the complex nature of tauopathies.

## Acknowledgements

We thank Dr. Peter Davies (Albert Einstein College of Medicine, USA), and Dr. J.E. Darnell, Jr. (Rockefeller University) for providing the materials (antibodies and plasmids) for this study. We thank Dr. Tiago Outerio for providing the materials (VN-Tau (P301L), Addgene plasmid #87634; Tau (P301L)-VC, Addgene plasmid #87633) for this study. We would also like to thank Seoul National University Hospital Brain Bank (SNUHBB; 2023-ER1005-01) and Pusan National University Hospital Brain Bank (PNUHBB; 2023-ER1002-01) for providing the samples of postmortem brain tissue and cerebrospinal fluid for this study.

## Author contributions

Yoon G and Jo C, Conceptualization; Yoon G and Park H, Data curation; Yoon G, Park H, Park JY, Choi JY, Kam MK, Formal analysis; Jo C and Kim H, Funding acquisition; Yoon G, Park H, Kam MK, Kim JH, Jo C, Investigation; Yoon G and Park H, Methodology; Jo C, Kim H, Kim JH, Project administration; Park SH, Kim EJ, Resources; Jo C, Supervision; Yoon G, Park H, Park JY, Kam MK, Choi JY, Validation; Yoon G, Park H, Jo C, Writing – original draft preparation; Johnson GV, Kim H, Kim JH, Koh YH, Writing – review & editing.

## Institutional review board statement

This study was approved by the Institutional Review Board of the Korea Disease Control and Prevention Agency (reference number: 2022-07-06-2C-A).

## Informed consent statement

All subjects who participated in the study provided their informed consent.

## Data availability statement

Data supporting the findings of this study can be obtained from the corresponding author upon request.

## Materials and Methods

The following procedures were conducted in accordance with established protocols outlined in the Online Methods section: ethical statement, human brain tissue, human cerebrospinal fluid (CSF), cell lines and culture, iPSC-derived cortical neuron and brain organoids, generation of stable cell lines, plasmid construction and subcloning, preparation of AAV particles, transfection and treatment, tau propagation analyses, cDNA microarray, mRNA-sequencing, bioinformatic analyses, exosome isolation, total membranes isolation, calcium staining, western blot analysis, human phosphorylated tau ELISA, immunoprecipitation, fluorescent imaging, and statistical analyses.

## Online Methods

### Ethic statement

Participants were identified from the Seoul National University Hospital Brain Bank (SNUHBB) and Pusan National University Hospital Brain Bank (PNUHBB), and written informed consent was obtained from each patient or their legal guardian prior to death. This study adhered to the guidelines approved by the Institutional Review Board of the Korea Disease Control and Prevention Agency (KDCA; IRB number: 2022-07-06-2C-A), which governs the use of human brain tissue and cerebrospinal fluids. All experimental procedures were conducted in accordance with established ethical standards and guidelines. This research was conducted in compliance with the Declaration of Helsinki and the ethical principles outlined by the KDCA. Additionally, the confidentiality and anonymity of participants were maintained throughout the study. The data used in this study were anonymized prior to analysis to maintain the privacy of the participants.

### Human brain tissue

The human brain tissues used in this investigation were sourced from SNUHBB and PNUHBB, where postmortem neuropathological examinations were conducted. Diagnoses of VD, CBD, PSP, PART, and AD were made according to the criteria stipulated by the National Institute on Aging and Alzheimer’s Association guidelines (NIA-AA). Brain tissues were collected following standard protocols and immediately frozen at -80°C for storage prior to analysis. Immunoblot analysis was employed to analyze the brain tissues following the protocols detailed in the western blot analysis section, below.

### Human cerebrospinal fluid

The human CSF used in this study was obtained from PNUHBB. Prior to inclusion, each participant provided informed consent in accordance with established guidelines and regulations. A neurologist specializing in neurodegenerative diseases conducted comprehensive evaluations, including clinical interviews and neurological examinations. The diagnoses of mild cognitive impairment and AD were diagnosed according to the NIA-AA core clinical criteria and modified Petersen criteria, respectively. CSF collection, involving nine subjects, adhered to established protocols and was conducted at the Research Institute for Convergence of Biomedical Science and Technology at the Pusan National University Yangsan Hospital. CSF samples were immediately frozen and stored at -80°C prior to analysis. The quantification of FAM84B levels was performed using immunoblot analysis following the procedures outlined in the western blot analysis section below.

### Cell lines and culture

Mouse Neuro-2A, human HEK293, and SH-SY5Y neuroblastoma cells were obtained from the American Type Culture Collection (ATCC). Neuro-2A and HEK293 cells were cultured in Dulbecco’s Modified Eagle medium (DMEM; Thermo Fisher Scientific, Waltham, MA, USA) supplemented with 10% fetal bovine serum (FBS; Merck Millipore, Burlington, MA, USA), 1 mM sodium pyruvate (Thermo Fisher Scientific), and 100 U/ml penicillin-streptomycin (Thermo Fisher Scientific). SH-SY5Y cells were cultured in DMEM/F12 containing 10% FBS and 100 U/ml penicillin-streptomycin. All cells were maintained at 37°C in a 5% CO2 atmosphere, and the medium was refreshed every two days. Sub-culturing was performed using prewarmed 1X PBS (Thermo Fisher Scientific) and 0.25% trypsin-EDTA (Thermo Fisher Scientific).

Neuronal differentiation of Neuro-2A and SH-SY5Y cells was initiated by supplementing the DMEM/F12 medium with 10 μM all-trans retinoic acid (Sigma-Aldrich, Burlington, MA, USA), 2% FBS, and 100 U/ml penicillin-streptomycin. The medium was renewed every two days until the cells reached a differentiated state by day 5.

Human induced pluripotent stem cells (hiPSCs) were cultured on vitronectin-coated plates (Gibco, Thermo Fisher Scientific) in TeSR-E8 medium (Stem Cell Technologies, Vancouver, CA). The culture medium was replaced daily. For subculturing, the cells were dissociated using EDTA (Gibco). The normal hiPSC lines hFSiPS3-1 and PB01-EiPS21 were procured from the Korea National Stem Cell Bank. The AD hiPSC lines UCSD234i-SAD2-3 and UCSD241i-APP2-3 were obtained from WiCell Research Institute Inc (Madison, WI, USA)^40, 41^.

### hiPSC-derived cortical neuron and brain organoids

Differentiation of hiPSC-derived cortical neurons and brain organoids was performed as previously described, with some modifications^42^. Briefly, hiPSCs were dissociated using Accutase® solution (Sigma-Aldrich). A total of 9,000 cells were cultured in AggreWell™ EB formation medium (Stem Cell Technologies) with 1X CloneR™ (Stem Cell Technologies) in Ultra Low Cluster 96-well plates (Costar, Corning, NY, USA). On day 2, the formed embryoid bodies (EBs) were transferred to a petri dish (SPL) and cultured in AggreWell™ EB formation medium supplemented with 2.5 µM Dorsomorphin dihydrochloride (Tocris Bioscience, Bristol, UK) and 10 µM SB431542 (Tocris), with daily medium changes. On day 7, EBs were cultured in neurobasal medium (Gibco) supplemented with 2% B-27 without vitamin A (Gibco), 1% GlutaMax (Gibco), 20 ng/ml EGF (R&D Systems, Minneapolis, MN, USA), and 20 ng/ml FGF-2 (PeproTech, Cranbury, NJ, USA), with daily medium changes.

On days 21–23, for cortical neuron differentiation, EBs were dissociated using 0.05% Trypsin-EDTA (Gibco) and plated on dishes coated with 50 µg/ml Poly-L-ornithine (Sigma-Aldrich) and 20 µg/ml Laminin (Sigma-Aldrich). Cells were cultured in DMEM/F12 medium (Gibco) supplemented with antibiotic-antimycotic (Gibco), 2% B-27 without vitamin A (Gibco), 1% N-2 (Gibco), 20 ng/ml brain-derived neurotrophic factor (BDNF; R&D Systems), and 20 ng/ml NT-3 (R&D Systems). The medium was changed every other day. On day 25, for brain organoid differentiation, EBs were cultured in neurobasal medium (Gibco) supplemented with 2% B-27 without vitamin A (Gibco), 1% GlutaMax (Gibco), 20 ng/ml BDNF (R&D Systems), and 20 ng/ml NT-3 (R&D Systems). The medium was changed every other day. Cortical neurons and brain organoids were cultured for at least 40 days.

### Generation of stable cell lines

To establish stable tau-expressing cell lines, HEK293 cells were transfected with pcDNA3.1-Tau, which carries a neomycin resistance gene, using Lipofectamine 3000 (Thermo Fisher Scientific). Following a 48-hour post-transfection period, neomycin selection was initiated with G418 (Gibco) at doses determined via a kill curve (1–5 μg/ml). Neomycin-resistant cells and individual cell colonies were subsequently isolated using cloning rings and protein expression levels were assessed using western blot analysis.

To generate stable FAM84B knockout cell lines, we used the FAM84B human gene knockout kit (OriGene, Rockville, MD, USA) following the manufacturer’s protocol. HEK293 cells were transfected with FAM84B gRNA vectors (or scrambled controls) and linear donor DNA (LoxP-EF1A-tGFP-P2A-Puro-LoxP) using Lipofectamine 3000 (Thermo Fisher Scientific). After 48 hours post-transfection, cells were passaged and cultured for 3 days, with subsequent passages every 3 days for a total of seven passages. Puromycin (Invitrogen, Thermo Fisher Scientific) selection was initiated at the 5^th^ or 7^th^ passage, with doses determined via a kill curve (1–10 μg/ml). Puromycin-resistant cells and individual cell colonies were isolated using cloning rings and analyzed by western blotting and genomic PCR to confirm gene knockdown and integration of the functional cassette.

### Plasmid construction and subcloning

A combination of plasmids was used to elucidate the role of FAM84B in tauopathy. These included: pcDNA3.1-Tau (Gail VW Johnson Lab), pCMV6-FAM84B-myc-DDK (OriGene #RC207996), pEGFP-C1-Tau (Gail VW Johnson Lab), pmCherry-C1-FAM84B (subcloning using *KpnI*/*EcoRI*), pcDNA3.1-Tau-p2a-eGFP (subcloning using *BamHI*/*XhoI*), pcDNA3.1-Tau(P301L)-VC (Addgene #87633), pcDNA3.1-VN-Tau(P301L) (Addgene #87634), pcDNA3.1-STAT3-WT (JE Darnell Jr Lab), and pcDNA3.1-STAT3-DN (JE Darnell Jr Lab). All subcloned plasmids were confirmed by sequencing and protein expression analysis and were used for transient transfection. The primer sequences used are listed in Supplementary Table 1.

Additionally, we used plasmids from the AAVpro Helper Free Systems (Takara Bio, Kusatsu, Japan) to investigate the role of FAM84B in iPSC-derived cortical neurons and brain organoids. These included: pAAV-FAM84B (subcloning using *EcoRI*/*BamHI*), pAAV-FAM84B-myc (subcloning using *EcoRI*/*BamHI*), and pAAV-Tau-myc-p2a-eGFP (subcloning using *EcoRI*/*XbaI*). All plasmids were confirmed by sequencing and used for AAV infection. The primer sequences used are listed in Supplementary Table 1.

### Preparation of AAV particles

AAV6 particles were produced by co-transfecting HEK293T cells (ATCC) with a triple plasmid system [pAAV, pRC6, and pHelper] provided by the AAVpro Helper Free system (Takara Bio). After 72 h, AAV particles were harvested from the transfected cells using an AAVpro purification kit (Takara Bio). Recombinant AAV vectors were titrated by measuring the expression levels of the proteins of interest using western blot analysis.

### Transfection and treatment

Human HEK293 and SH-SY5Y cells were transfected with pcDNA3.1-Tau, pcDNA3.1-Tau(P301L)-VC, pcDNA3.1-VN-Tau(P301L), pCMV6-FAM84B-myc-DDK, pcDNA3.1-STAT3-WT, or pcDNA3.1-STAT3-DN using Lipofectamine 3000 (Thermo Fisher Scientific) following the manufacturer’s protocol. All analyses were conducted at 48 h post-transfection.

For assessing neuroinflammation-mediated FAM84B expression, human SH-SY5Y cells and iPSC-derived cortical neurons were treated with sterilized distilled water (control vehicle), 50 ng/μl IL-6, or 50 ng/μl TNF-α, either separately or in combination. Analyses were performed at 48 h post-treatment.

To inhibit RYR3-mediated cytosolic calcium regulation, human HEK293 and SH-SY5Y cells, along with iPSC-derived cortical neurons, were treated with DMSO (control vehicle) or 10 μM Dantrolene. All analyses were conducted at 24 h post-treatment.

To inhibit clathrin-mediated endocytosis, human SH-SY5Y cells were treated with DMSO (control vehicle) or 10 μM Pitstop2. Analyses were performed at 24 h post-treatment.

### Tau propagation analyses

Two distinct methodologies were employed to validate FAM84B-mediated tau propagation.

To visualize tau propagation upon exposure to conditioned media, human HEK293 and SH-SY5Y cells were transfected with pcDNA3.1-Tau or pCMV6-FAM84B-myc-DDK for 48 h. Subsequently, the conditioned medium from these cells was transferred to normal HEK293 and SH-SY5Y cells for an additional 24 h.

To visualize the transmission of tau from donor to recipient cells, human HEK293 and SH-SY5Y cells were transfected with pcDNA3.1-Tau-p2a-eGFP and pCMV6-FAM84B-myc-DDK for 48 h. Additionally, iPSC-derived cortical neurons and brain organoids were infected with AAV6 particles containing FAM84B-Myc for 4 days and analyzed after 6 days of treatment in the growth medium.

The analysis of tau propagation assessed the proportion of tau spread events, characterized by an increase in the number of recipient cells (green fluorescent protein negative/red signal positive) relative to the total number of donor cells (green fluorescent protein positive and/or red signal negative).

### cDNA microarray

Total RNA was extracted from cells expressing either relatively low or high levels of tau protein using TRIzol reagent (Invitrogen) according to the manufacturer’s protocol. The RNA purity and integrity were assessed using an ND-1000 Spectrophotometer (NanoDrop Technologies, Wilmington, DE, USA) and an Agilent 2100 Bioanalyzer (Agilent Technologies, Inc., Santa Clara, CA, USA). Total RNA was amplified and purified using the TargetAmp-Nano Labeling Kit for Illumina Expression BeadChip (EPICENTRE; Illumina Inc., San Diego, CA, USA) to generate biotinylated cRNA according to the manufacturer’s instructions.

Briefly, 500 ng of total RNA was reverse-transcribed into cDNA using T7 oligo(dT) primers. The subsequent steps included second-strand cDNA synthesis, in vitro transcription, and biotin-NTP labeling. After purification, the cRNA was quantified using an ND-1000 Spectrophotometer. Each 750 ng of labeled cRNA sample was hybridized to a Human HT-12 v4.0 Expression BeadChip for 17 hours at 58°C, following Illumina’s instructions. Array signal detection was performed using Amersham Fluorolink Streptavidin-Cy3 (GE Healthcare Bio-Sciences, Piscataway, NJ, USA) according to the bead array manual, and the arrays were scanned using an Illumina BeadArray Reader confocal scanner.

Quality control checks were performed visually on the internal controls and raw scanned data. Raw data were extracted using Illumina GenomeStudio v2011.1 software (Gene Expression Module v1.9.0), followed by logarithmic transformation and quantile normalization of the array probes. The statistical significance of the expression data was assessed using the LPE test and fold change analysis, with the false discovery rate (FDR) controlled via the adjustment of p-values using the Benjamini-Hochberg algorithm. Hierarchical cluster analysis of differentially expressed gene sets was conducted using complete linkage and Euclidean distance as similarity measures.

### mRNA sequencing

mRNA was extracted from the negative control, FAM84B knockout, FAM84B overexpression, and Tau/FAM84B co-overexpressing HEK293 and SH-SY5Y cells using TRIzol Reagent (Thermo Fisher Scientific), according to the manufacturer’s instructions. DNase I (Takara Bio) was added and incubated for one hour to remove any residual DNA. The RNA concentration was quantified using an ND-1000 spectrophotometer (Thermo Fisher Scientific), and RNA integrity was assessed using a 2100 Expert Bioanalyzer (Agilent Technologies, Inc.).

rRNA was depleted using a Ribo-Zero Gold rRNA Removal Kit (Illumina, Inc.). Following rRNA depletion, RNA-sequencing libraries were prepared using the TruSeq Stranded mRNA Kit (Illumina, Inc.) and sequenced on a NovaSeq 6000 (Illumina, Inc.) with 2×100 bp paired-end reads.

RNA-Seq data analysis involved mapping the reads to the reference genome (mouse mm10, human hg19) using Tophat (v2.0.13), followed by differential expression analysis using Cuffdiff (v2.2.0). Library normalization and dispersion estimation were performed using geometric and pooled methods depending on the availability of replicates. CummeRbund (v2.8.2) was used for data visualization. DEGs were identified from the "gene_exp.diff" output of Cuffdiff, filtering for genes with a "OK" status and a fold change exceeding 2.

### Bioinformatic analyses

For the gene ontology (GO) analysis of the selected genes from the mRNA-seq data, we utilized the Gene Ontology Resource (http://geneontology.org/). Altered genes were analyzed by selecting their "Cellular compartment" terms. The PANTHER classification system and Fisher’s exact test were employed, followed by the calculation of the false discovery rate (FDR) and fold enrichment. The selected genes and GO terms utilized are listed in Supplementary Table 2.

To predict the transcription factors and chromatin regulators involved in FAM84B expression, we used the ChIP-seq database (https://maayanlab.cloud/Harmonizome/), which includes the ENCODE and ChEA3 systems. This list is included in Supplementary Table 2.

To ascertain the binding regions of the predicted transcription factors to the FAM84B promoter, genomic information was sourced from the human genome (GRCh37/hg19) available on the UCSC Genome Browser (https://genome.ucsc.edu/). We then accessed public data from ENCODE ChIP-seq, GeneHancer regulatory elements and genes, and the FANTOM5 enhancer in the UCSC Genome Browser. We aligned and displayed the predicted transcription factors bound to the enhancer regions of FAM84B.

### Exosome isolation

The membrane affinity-based exosome isolation process began with filtering the conditioned media through a 0.45 μm syringe filter (Pall Corporation, New York, NY, USA). The filtered media were then mixed with a binding buffer (buffer XBP) and loaded onto the membrane spin column of the ExoEasy Maxi Kit (Qiagen, Venlo, the Netherlands). After centrifugation at 500×g for 3 min, vesicles were captured within the membrane, whereas excess fluid and proteins were removed. The bound vesicle fraction was purified by adding washing buffer (buffer XWP) to the membrane, followed by centrifugation at 5,000×g for 5 min.

After replacing the collection tube, the elution buffer (buffer XE) was added to the column and incubated for 1 min at room temperature. Centrifugation at 500×g for 5 min facilitated the elution of the exosome fraction. To enhance exosome recovery, the eluate was reapplied to the column, incubated for 1 min at room temperature, and centrifuged at 5,000×g for 5 min. Finally, the exosome fraction was collected in a final volume of 200 μl of the provided elution buffer.

### Total membrane isolation

The total membrane fraction was determined using a Plasma Membrane Protein Extraction Kit (Abcam, Cambridge, UK). Initially, the cells were harvested by centrifugation at 600×g for 5 min, washed with ice-cold 1× phosphate-buffered saline (PBS), and suspended in the homogenization buffer mix provided by the kit. After homogenization, the resulting homogenate was centrifuged at 700×g for 10 min to remove cellular debris, and the supernatant was collected to isolate total cellular membrane proteins. After centrifugation at 10,000g for 30 min, the cytosolic fraction remaining in the supernatant was separated from the total membrane fraction contained within the pellet.

### Calcium staining

Cytosolic calcium levels in HEK293 and SH-SY5Y cells were assessed using Fluo-4 AM solution (Invitrogen) according to the manufacturer’s protocol. A 3 mM Fluo-4 AM/DMSO stock solution was diluted to a 3 µM working solution in the respective culture medium. Cells were incubated with the Fluo-4 AM working solution for 20 minutes at 37°C in a CO2 incubator. After incubation, the cells were washed with indicator-free medium to remove non-specifically associated dyes and then incubated with indicator-free medium for an additional 30 min to ensure complete de-esterification of intracellular AM esters. Calcium imaging and cellular morphology analyses were performed using the EVOC M5000 microscope (Invitrogen).

### Western blot analysis

Western blot analysis was performed according to established protocols. Cells and tissues were lysed in ice-cold RIPA buffer (GenDEPOT, Altair, TX, USA) for 10 min on ice, and protein concentrations were determined using a BCA assay kit (Thermo Fisher Scientific) following the manufacturer’s instructions. Protein samples (15–25 μg) were separated on 8–12% pre-made SDS-polyacrylamide gels (Thermo Fisher Scientific) and transferred onto polyvinylidene difluoride (Merck Millipore) membranes using absolute methanol (Thermo Fisher Scientific). The membranes were blocked in a solution containing 5% bovine serum albumin (GenDEPOT) and skim milk (Cell Signaling Technology, Danvers, MA, USA) for one hour at room temperature to enhance the detection of the phosphorylated and native forms of the proteins. Following blocking, membranes were incubated overnight at 4°C with primary antibodies (1:1,000 dilution), including: PHF-1 (Peter Davies Lab), Tau5 (Gail VW Johnson Lab), Tau (Dako, Agilent Technologies, Inc., 0024), Myc-tag (Cell Signaling Technology, 2276), FAM84B (OriGene, TA501992; Proteintech Group, Inc., Rosemont, IL, USA, 18421-1-AP), Alix (Cell Signaling Technology, 2171), Caveolin-1 (Cell Signaling Technology, 3267), RYR3 (Alomone Labs, Jerusalem, Israel, ARR003), Clathrin-heavy chain (CHC, Cell Signaling Technology, 4796), EEA1 (Cell Signaling Technology, 3288), Rab5 (Cell Signaling Technology, 3547), Rab7 (Cell Signaling Technology, 9367), Rab9 (Cell Signaling Technology, 5118), Rab11 (Cell Signaling Technology, 5589), pSTAT3 (Y705) (Cell Signaling Technology, 9145), STAT3 (Cell Signaling Technology, 9139), p62 (Cell Signaling Technology, 8025), LC3 (Cell Signaling Technology, 3868), HSP70 (Cell Signaling Technology, 4876), LAMP-1 (BD Biosciences, Franklin Lakes, NJ, USA, 611042), LAMP-2 (Sigma-Aldrich, L0668), and beta-actin (Merck Millipore, MAB1501).

After incubation with the primary antibody, the membranes were probed with the corresponding horseradish peroxidase-conjugated secondary antibody (1:10,000 dilution; BioLegend, San Diego, CA, USA) for one hour at room temperature. Protein bands were visualized using an ECL solution (Thermo Fisher Scientific) and imaged using a ChemiDocTM Imaging System (Bio-Rad, Hercules, CA, USA). Quantification of protein expression levels was performed using Image J software (V1.53c, NIH, Bethesda, MD, USA), with normalization against beta-actin and respective native proteins.

### Human phosphorylated tau ELISA

Conditioned media from the experimental samples were analyzed using a human phosphorylated tau (T181/S396) ELISA kit (Invitrogen). Standards were prepared by diluting the supplied standard with "standard dilution buffer" to a concentration of 2,000 pg/mL, followed by serial dilutions to achieve concentrations ranging from 31.25 to 1,000 pg/mL.

Duplicate wells received 100 μL of standards, whereas the remaining wells were treated with 50 μL of standard dilution buffer and 50 μL of the sample in duplicate.

After incubating the ELISA plates for 2 h at room temperature on a rotary shaker, they were washed four times. Subsequently, 100 μL of the corresponding anti-p-tau detection antibody was added to each well and incubated for 1 h, followed by four washes. The anti-rabbit HRP concentrate was diluted 100-fold and 100 μL was added to each well, followed by a 30-minute incubation. Subsequently, 100 μL of stabilized chromogen was added to each well, and the plate was incubated at room temperature in darkness for 20 minutes on the rotary shaker. The reaction was stopped with 100 μL of the provided "stop" solution, and the plate was read at 450 nm using a SPECTRA MAX 190 plate reader (Molecular Devices, San Jose, CA, USA).

### Immunoprecipitation

Cells were lysed in NP-40 lysis buffer (0.5% NP-40, 10 mM Tris–HCl [pH 8.0], 150 mM NaCl, 10 mM sodium pyrophosphate, and 1 mM EDTA) supplemented with 1 mM NaF, 1 mM Na3VO4, and protease inhibitor cocktail. Equal amounts of lysates were then incubated with 2 μg of antibodies pre-conjugated with sheep anti-mouse magnetic beads (DYNAL, Invitrogen) for 3 hours on a rotational shaker at 4°C. After overnight incubation at 4°C, the beads were washed thrice with NETN wash buffer (0.1% NP-40, 50 mM Tris–HCl [pH 8.0], 150 mM NaCl, 1 mM EDTA). The beads were then boiled in 1× SDS-sample loading buffer and 1× reducing agent at 95°C for 5 min. Proteins were subsequently separated by SDS-PAGE and immunoblotted under reducing and non-reducing conditions as previously described.

### Fluorescent imaging

The HEK293 and SH-SY5Y cells were fixed in 2% paraformaldehyde (Sigma-Aldrich) for 15 min, followed by overnight incubation with primary antibodies, including PHF-1, Tau5, Tau (Dako), and FAM84B in gelatin blocking buffer [0.1% gelatin (Sigma-Aldrich), 0.3% Triton X-100 (Thermo Fisher Scientific), 16 mM sodium phosphate (Sigma-Aldrich), 450 mM NaCl (Sigma-Aldrich), pH 7.4] at 4°C. After three washes with 1× PBS, the cells were incubated with Alexa 405/488/564-conjugated secondary antibodies (Invitrogen) for 2 h at room temperature. Following additional washes with 1× PBS, the cells were counterstained and mounted using the ProLong Gold Antifade Reagent (Thermo Fisher Scientific).

To detect fluorescent tagged proteins, including GFP-tau and mCherry-FAM84B, cells were fixed in 2% paraformaldehyde for 15 min. After three washes with 1× PBS, the cells were counterstained and mounted using the ProLong Gold Antifade Reagent.

hiPSC-derived cortical neurons were fixed in 4% paraformaldehyde (Biosesang, Seongnam, Korea) for 20 min, permeabilized in phosphate-buffered saline (PBS; GeneDEPOT) containing 0.25% Triton-X (Sigma-Aldrich) for 10 min, and blocked with 5% bovine serum albumin (BSA; Sigma-Aldrich) for 30 min. Cells were incubated overnight at 4°C with primary antibodies, including PHF-1 (Peter Davies Lab), Tau (Santa Cruz Biotechnology, Dallas, TX, USA, sc-32274), and FAM84B (OriGene, TA501992; Proteintech, 18421-1-AP). Subsequently, the cells were incubated with Alexa Fluor 488/594-conjugated secondary antibodies (Invitrogen) for one hour at room temperature. Nuclei were counterstained with 4’,6-diamidino-2-phenylindole (DAPI; Thermo Fisher Scientific) and mounted using Faramount Aqueous Mounting Medium (Dako).

Brain organoids were fixed in 4% paraformaldehyde (Biosesang) for one hour, washed with PBS, embedded in O.C.T (Tissue-Tek, Sakura Finetek USA, Inc., Torrance, CA, USA), and cryosectioned into 10-μm-thick slices. Immunocytochemistry was performed on the slices using the method described for cortical neurons.

Fluorescent images were captured using an EVOC M5000 microscope (Invitrogen), FV3000 microscope (Olympus, Tokyo, Japan), and LSM880 microscope (Carl Zeiss AG, Oberkochen, Germany). Colocalization analysis was performed using Pearson’s correlation test with the Coloc2 plugin in ImageJ (V1.53c, NIH).

### Statistical analyses

Data are presented as mean ± standard error of the mean unless specified otherwise. The experimental samples were randomly assigned to control or experimental groups, and the investigators were blinded to the experimental conditions. None of the samples were excluded from the analyses. For the in vitro, human brain tissue, and cerebrospinal fluid experiments group sample sizes were typically set to a minimum of three to ensure adequate statistical power. Normal distribution and homogeneity of variance within each comparison group were assessed prior to statistical analysis. Statistical comparisons between control and experimental samples were performed using an unpaired two-tailed *t*-test with Welch’s correction for unequal variances or ordinary two-way analysis of variance, as appropriate. Statistical significance was set at p < 0.05.

## References

1. Ritchie, C.W., Terrera, G.M. & Quinn, T.J. Dementia trials and dementia tribulations: methodological and analytical challenges in dementia research. Alzheimers Res Ther 7, 31 (2015).

2. Giacobini, E. & Gold, G. Alzheimer disease therapy--moving from amyloid-beta to tau. Nat Rev Neurol 9, 677–686 (2013).

3. Haass, C. & Selkoe, D. If amyloid drives Alzheimer disease, why have anti-amyloid therapies not yet slowed cognitive decline? PLoS Biol 20, e3001694 (2022).

4. Reitz, C., Pericak-Vance, M.A., Foroud, T. & Mayeux, R. A global view of the genetic basis of Alzheimer disease. Nat Rev Neurol 19, 261–277 (2023).

5. Zhang, Y., Wu, K.M., Yang, L., Dong, Q. & Yu, J.T. Tauopathies: new perspectives and challenges. Mol Neurodegener 17, 28 (2022).

6. Cummings, J.L., et al. The therapeutic landscape of tauopathies: challenges and prospects. Alzheimers Res Ther 15, 168 (2023).

7. Chen, Y. & Yu, Y. Tau and neuroinflammation in Alzheimer’s disease: interplay mechanisms and clinical translation. J Neuroinflammation 20, 165 (2023).

8. Cao, J., Hou, J., Ping, J. & Cai, D. Advances in developing novel therapeutic strategies for Alzheimer’s disease. Mol Neurodegener 13, 64 (2018).

9. Kapoor, M. & Chinnathambi, S. TGF-beta1 signalling in Alzheimer’s pathology and cytoskeletal reorganization: a specialized Tau perspective. J Neuroinflammation 20, 72 (2023).

10. Mueller, R.L., et al. Tau: A Signaling Hub Protein. Front Mol Neurosci 14, 647054 (2021).

11. Wei, Y., Liu, M. & Wang, D. The propagation mechanisms of extracellular tau in Alzheimer’s disease. J Neurol 269, 1164–1181 (2022).

12. Wu, J.W., et al. Neuronal activity enhances tau propagation and tau pathology in vivo. Nat Neurosci 19, 1085–1092 (2016).

13. Asai, H., et al. Depletion of microglia and inhibition of exosome synthesis halt tau propagation. Nat Neurosci 18, 1584–1593 (2015).

14. Rauch, J.N., et al. LRP1 is a master regulator of tau uptake and spread. Nature 580, 381–385 (2020).

15. van der Kant, R., Goldstein, L.S.B. & Ossenkoppele, R. Amyloid-beta-independent regulators of tau pathology in Alzheimer disease. Nat Rev Neurosci 21, 21–35 (2020).

16. Ljubenkov, P.A. & Rabinovici, G.D. Silencing tau to treat early Alzheimer’s disease. Nat Med 29, 1320–1321 (2023).

17. Sonnleitner, A., Conti, A., Bertocchini, F., Schindler, H. & Sorrentino, V. Functional properties of the ryanodine receptor type 3 (RyR3) Ca2+ release channel. EMBO J 17, 2790–2798 (1998).

18. Becherer, U., Moser, T., Stuhmer, W. & Oheim, M. Calcium regulates exocytosis at the level of single vesicles. Nat Neurosci 6, 846–853 (2003).

19. Hunter, C.A. & Jones, S.A. Corrigendum: IL-6 as a keystone cytokine in health and disease. Nat Immunol 18, 1271 (2017).

20. Arriagada, P.V., Growdon, J.H., Hedley-Whyte, E.T. & Hyman, B.T. Neurofibrillary tangles but not senile plaques parallel duration and severity of Alzheimer’s disease. Neurology 42, 631–639 (1992).

21. Xia, C., et al. Association of In Vivo [18F]AV-1451 Tau PET Imaging Results With Cortical Atrophy and Symptoms in Typical and Atypical Alzheimer Disease. JAMA Neurol 74, 427–436 (2017).

22. Braak, H. & Braak, E. Neuropathological stageing of Alzheimer-related changes. Acta Neuropathol 82, 239–259 (1991).

23. Ransohoff, R.M. How neuroinflammation contributes to neurodegeneration. Science 353, 777–783 (2016).

24. Ising, C., et al. NLRP3 inflammasome activation drives tau pathology. Nature 575, 669–673 (2019).

25. Heneka, M.T., et al. NLRP3 is activated in Alzheimer’s disease and contributes to pathology in APP/PS1 mice. Nature 493, 674–678 (2013).

26. Maphis, N., et al. Reactive microglia drive tau pathology and contribute to the spreading of pathological tau in the brain. Brain 138, 1738–1755 (2015).

27. Wong, N., et al. Upregulation of FAM84B during prostate cancer progression. Oncotarget 8, 19218–19235 (2017).

28. d’Errico, P. & Meyer-Luehmann, M. Mechanisms of Pathogenic Tau and Abeta Protein Spreading in Alzheimer’s Disease. Front Aging Neurosci 12, 265 (2020).

29. Huang, Y., et al. An in vitro vesicle formation assay reveals cargo clients and factors that mediate vesicular trafficking. Proc Natl Acad Sci U S A 118 (2021).

30. Samudra, N., Lane-Donovan, C., VandeVrede, L. & Boxer, A.L. Tau pathology in neurodegenerative disease: disease mechanisms and therapeutic avenues. J Clin Invest 133 (2023).

31. VandeVrede, L., Ljubenkov, P.A., Rojas, J.C., Welch, A.E. & Boxer, A.L. Four-Repeat Tauopathies: Current Management and Future Treatments. Neurotherapeutics 17, 1563–1581 (2020).

32. Chung, D.C., Roemer, S., Petrucelli, L. & Dickson, D.W. Cellular and pathological heterogeneity of primary tauopathies. Mol Neurodegener 16, 57 (2021).

33. Li, Q.S. & De Muynck, L. Differentially expressed genes in Alzheimer’s disease highlighting the roles of microglia genes including OLR1 and astrocyte gene CDK2AP1. Brain Behav Immun Health 13, 100227 (2021).

34. Creekmore, B.C., Watanabe, R. & Lee, E.B. Neurodegenerative Disease Tauopathies. Annu Rev Pathol 19, 345–370 (2024).

35. Nixon, R.A. The role of autophagy in neurodegenerative disease. Nat Med 19, 983–997 (2013).

36. Min, S.W., et al. Acetylation of tau inhibits its degradation and contributes to tauopathy. Neuron 67, 953–966 (2010).

37. Gallo, D., Ruiz, A. & Sanchez-Juan, P. Genetic Architecture of Primary Tauopathies. Neuroscience 518, 27–37 (2023).

38. Wyss-Coray, T. Inflammation in Alzheimer disease: driving force, bystander or beneficial response? Nat Med 12, 1005–1015 (2006).

39. Heneka, M.T., et al. Neuroinflammation in Alzheimer’s disease. Lancet Neurol 14, 388–405 (2015).

40. Uhm, K.O., et al. Generation of human induced pluripotent stem cell lines from human dermal fibroblasts using a non-integration system. Stem Cell Res 21, 13–15 (2017).

41. Sam Im, Y., Hoon Yoo, D., Kim, H.E., Young Oh, J. & Kim, Y.O. Generation of integration-free induced pluripotent stem cell line (KSCBi017-A) from peripheral blood mononuclear cells of a healthy male individual. Stem Cell Res 65, 102965 (2022).

42. Pasca, A.M., et al. Functional cortical neurons and astrocytes from human pluripotent stem cells in 3D culture. Nat Methods 12, 671–678 (2015).

