## Supplementary material for "FAM84B facilitates tau propagation via RYR3-mediated exocytosis in response to neuroinflammation": Supplemenatary file

**Supplementary Figure 1**

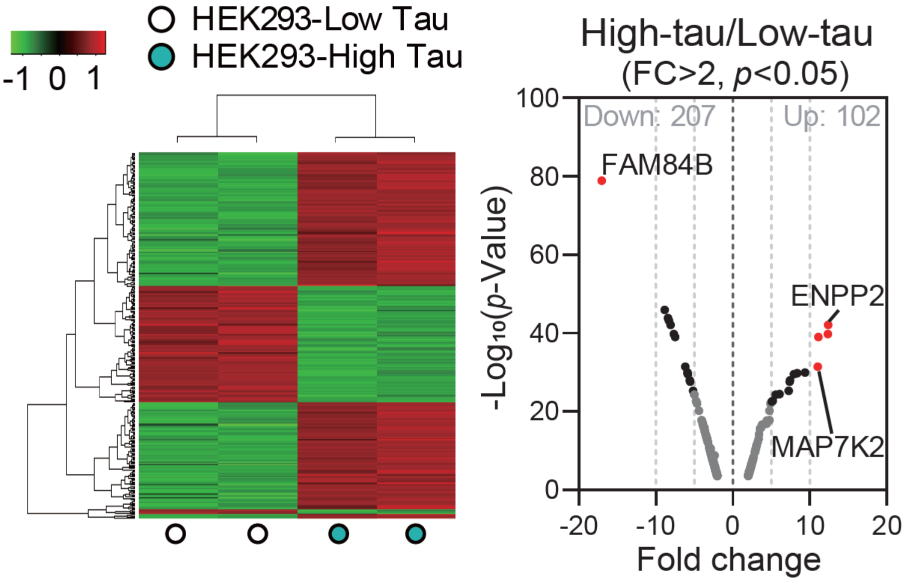

**Supplementary Figure 2**

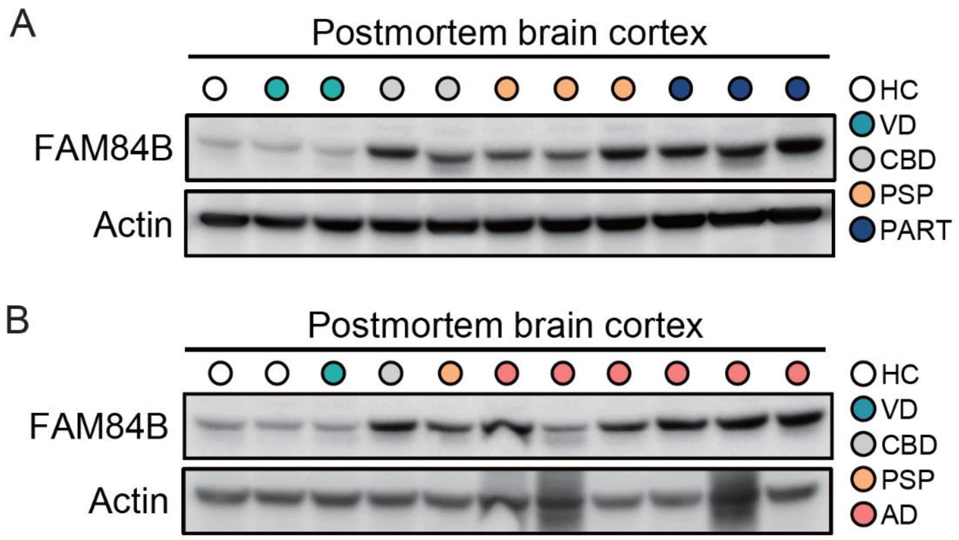

#### Supplementary Figure 3

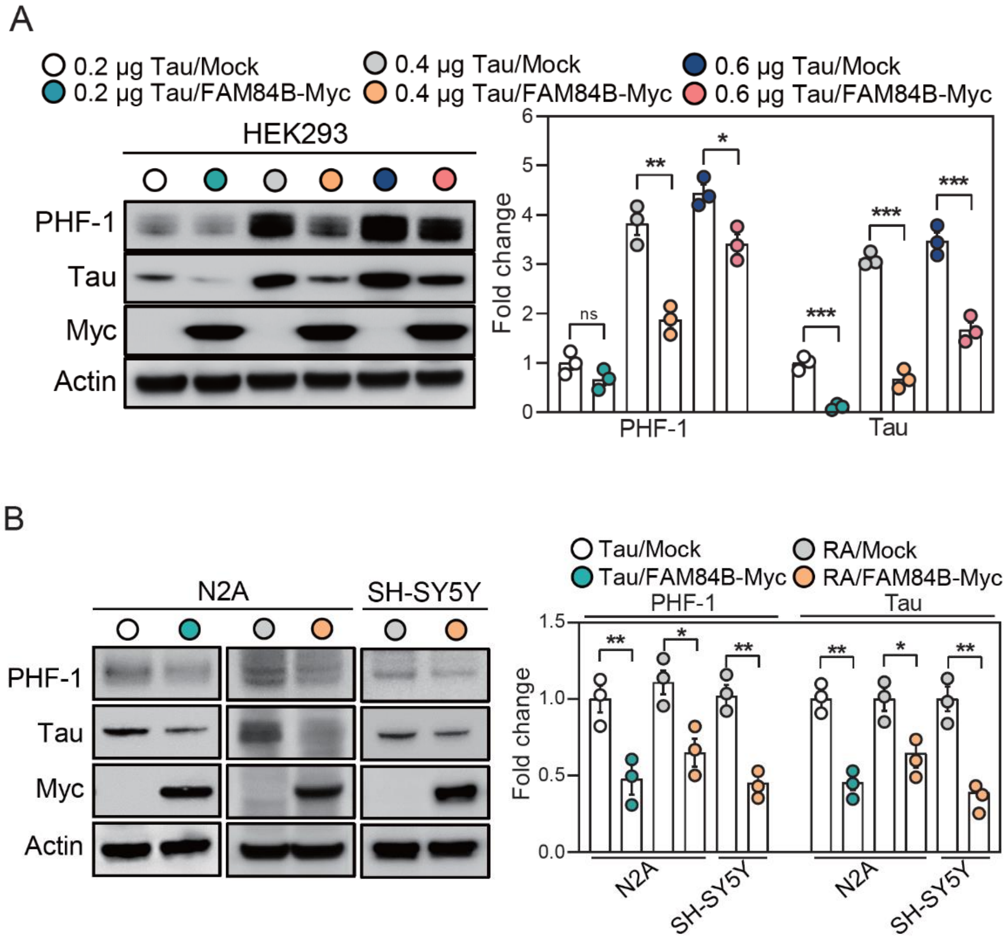

**Supplementary Figure 4**

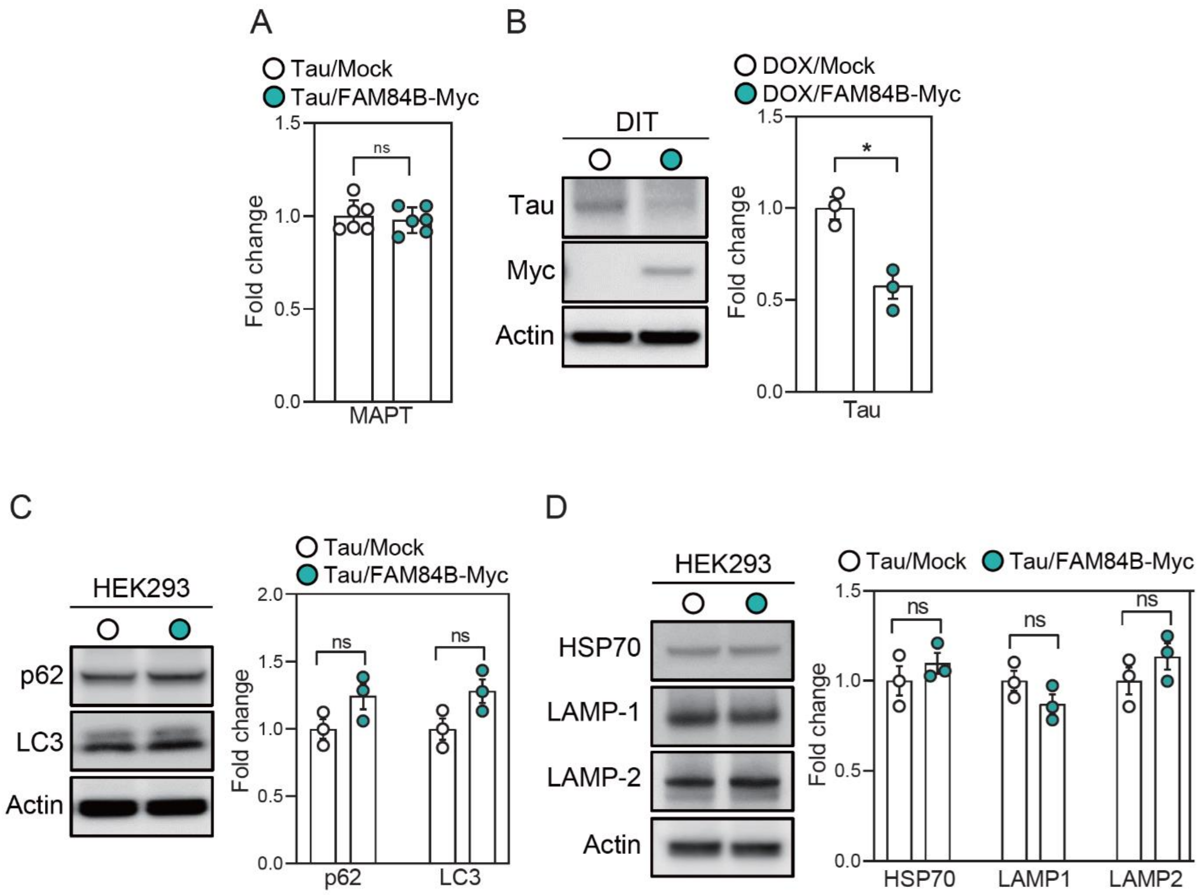

**Supplementary Figure 5**

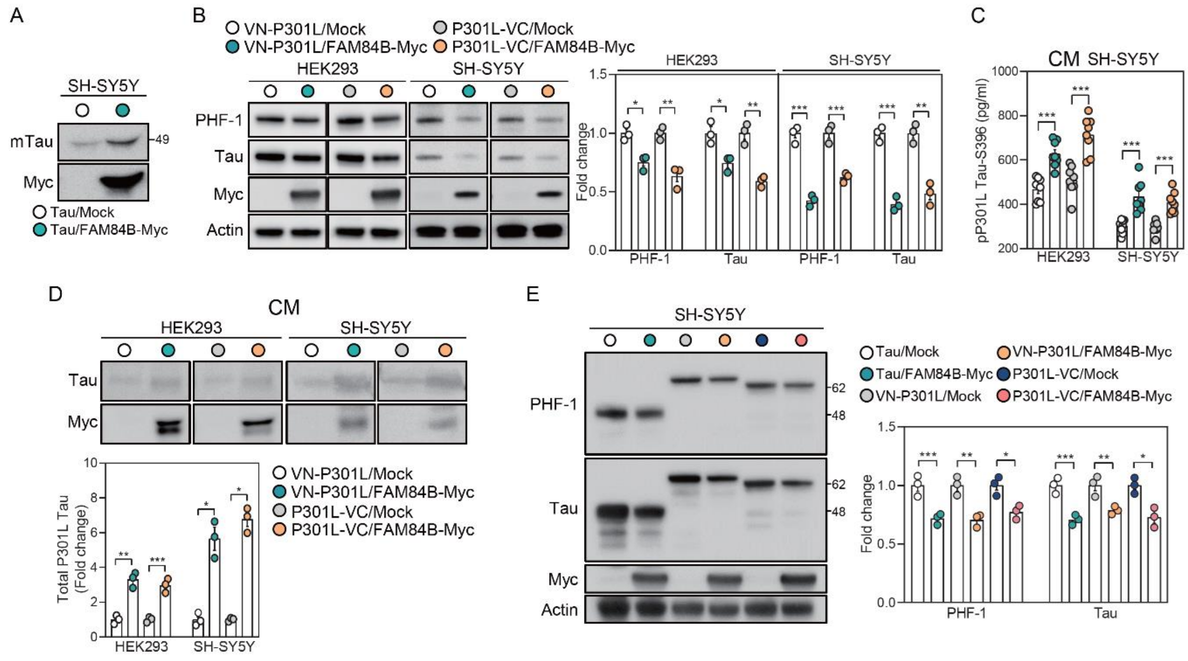

**Supplementary Figure 6**

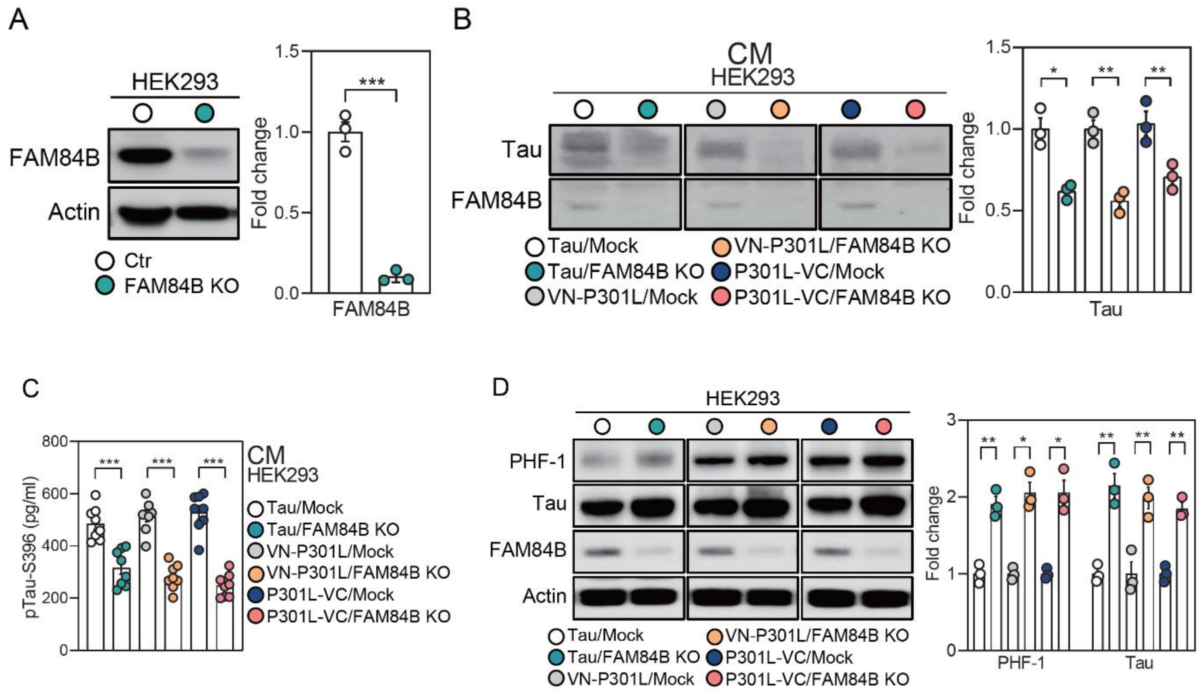

**Supplementary Figure 7**

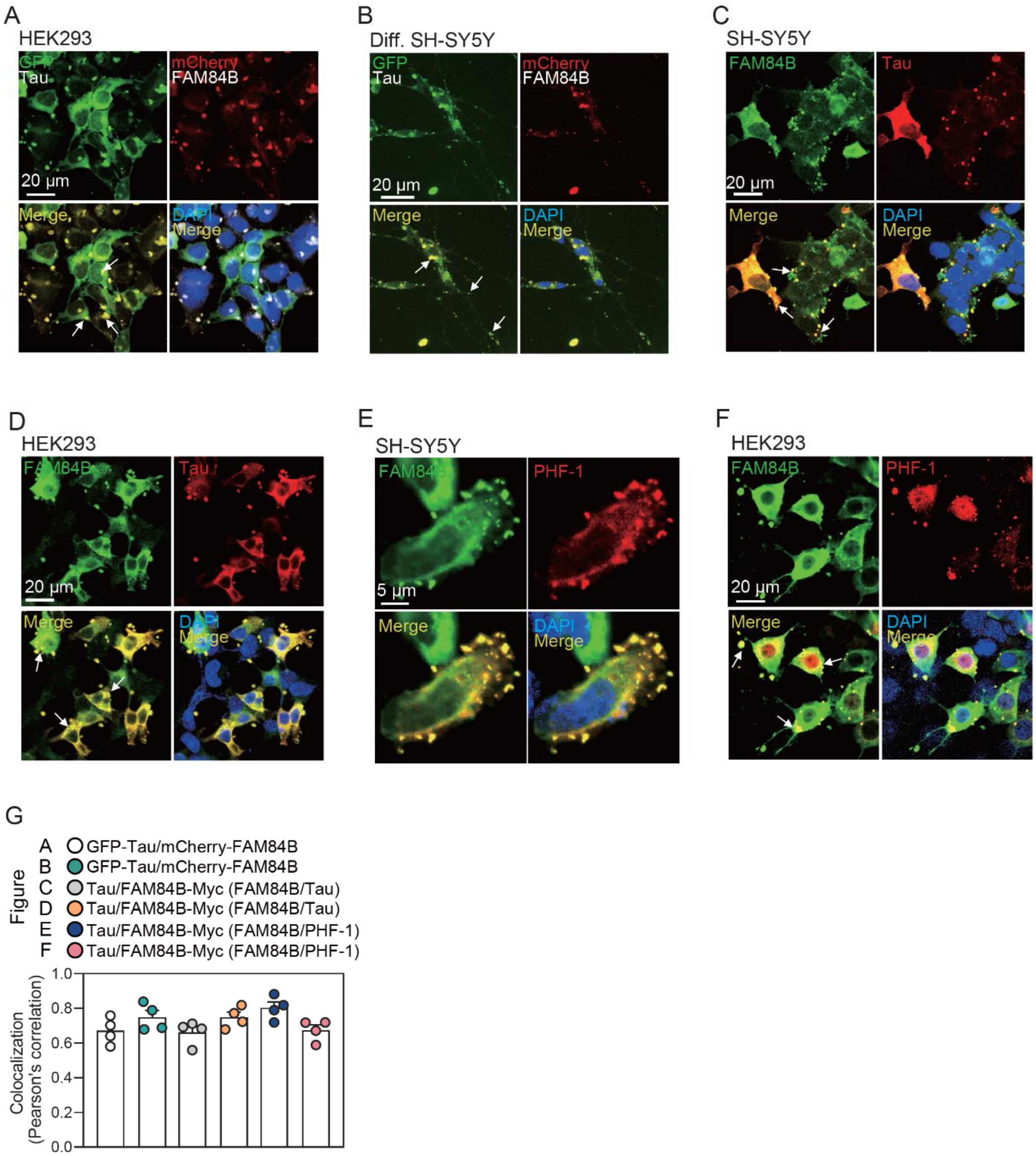

**Supplementary Figure 8**

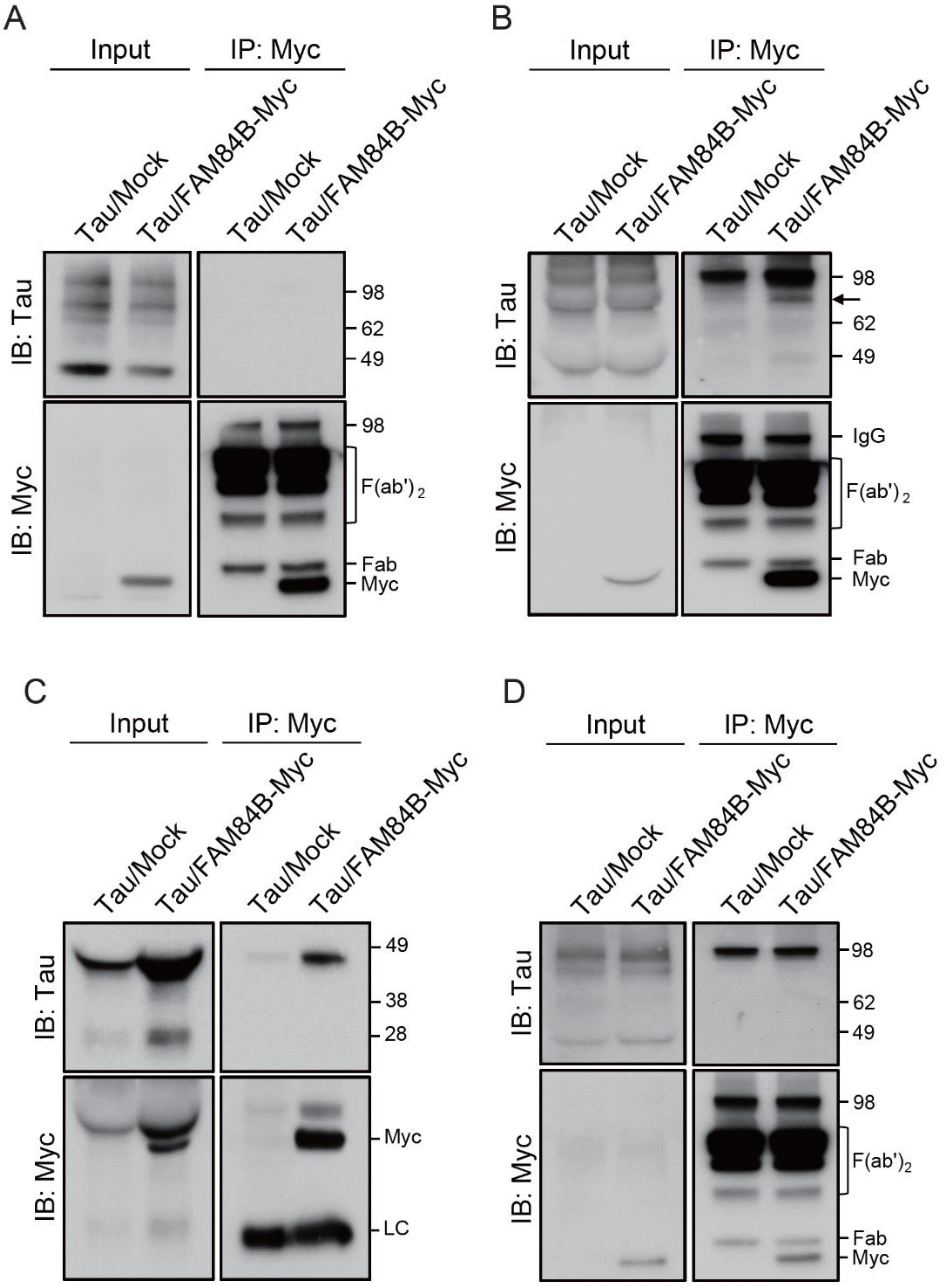

**Supplementary Figure 9**

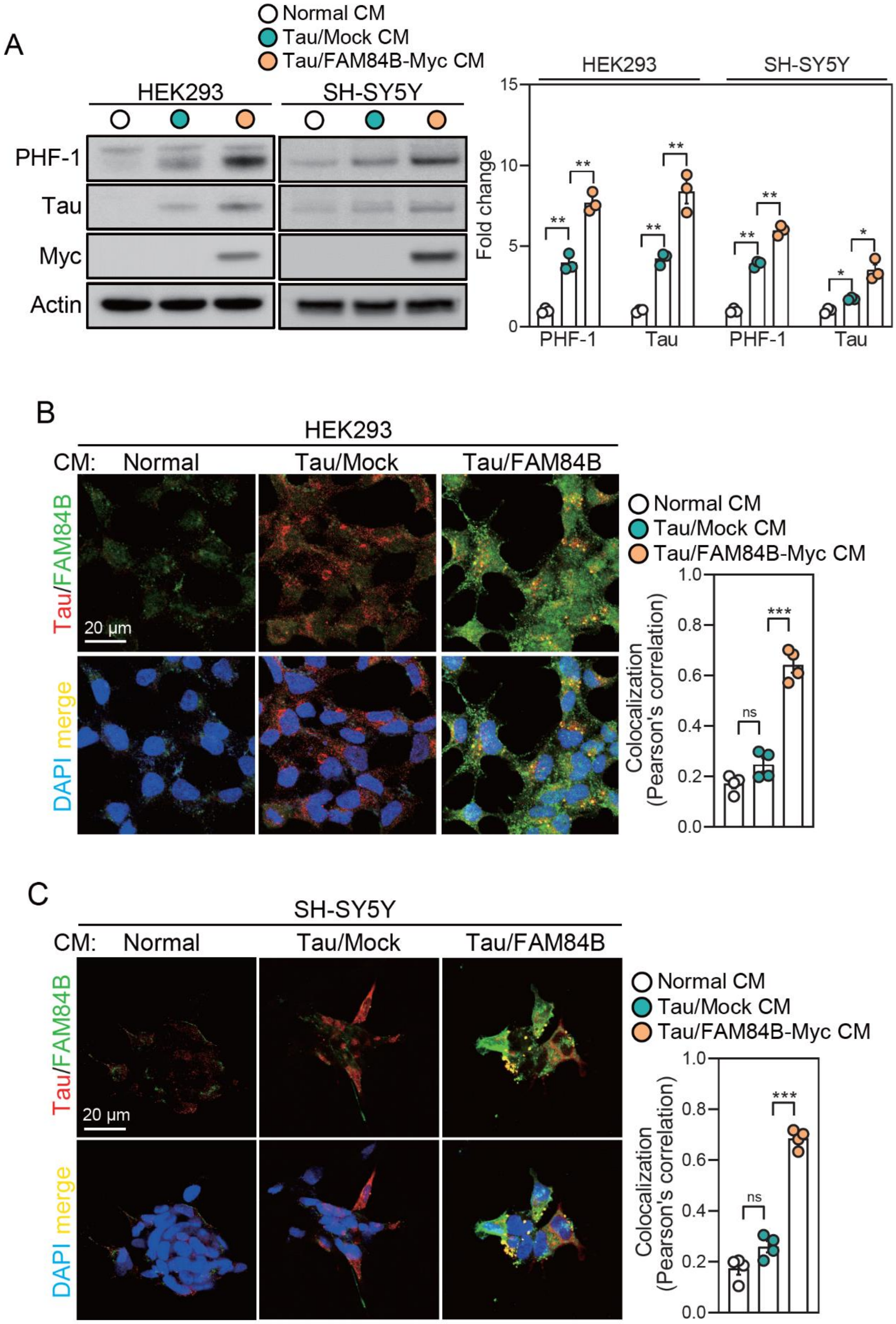

**Supplementary Figure 10**

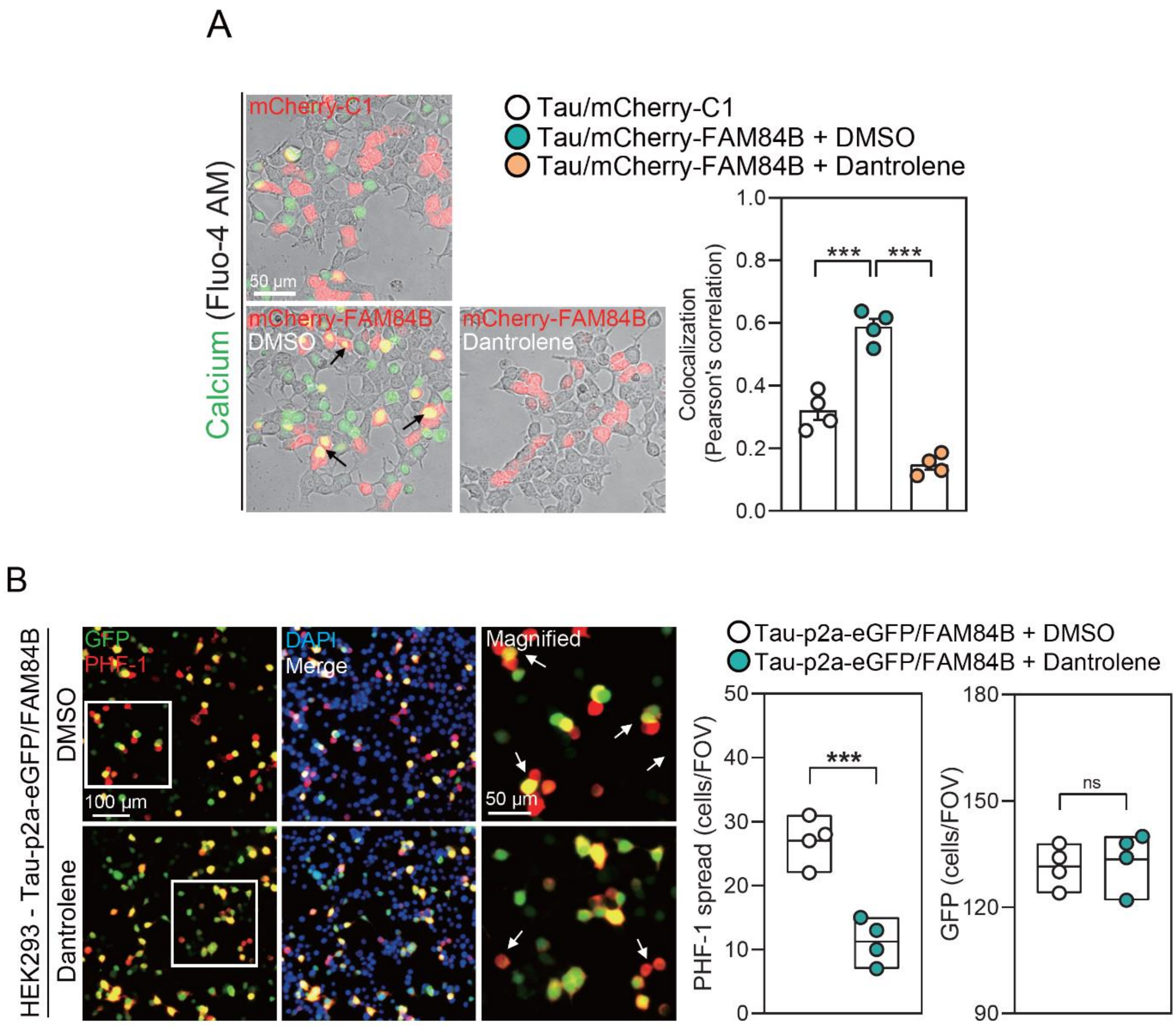

**Supplementary Figure 11**

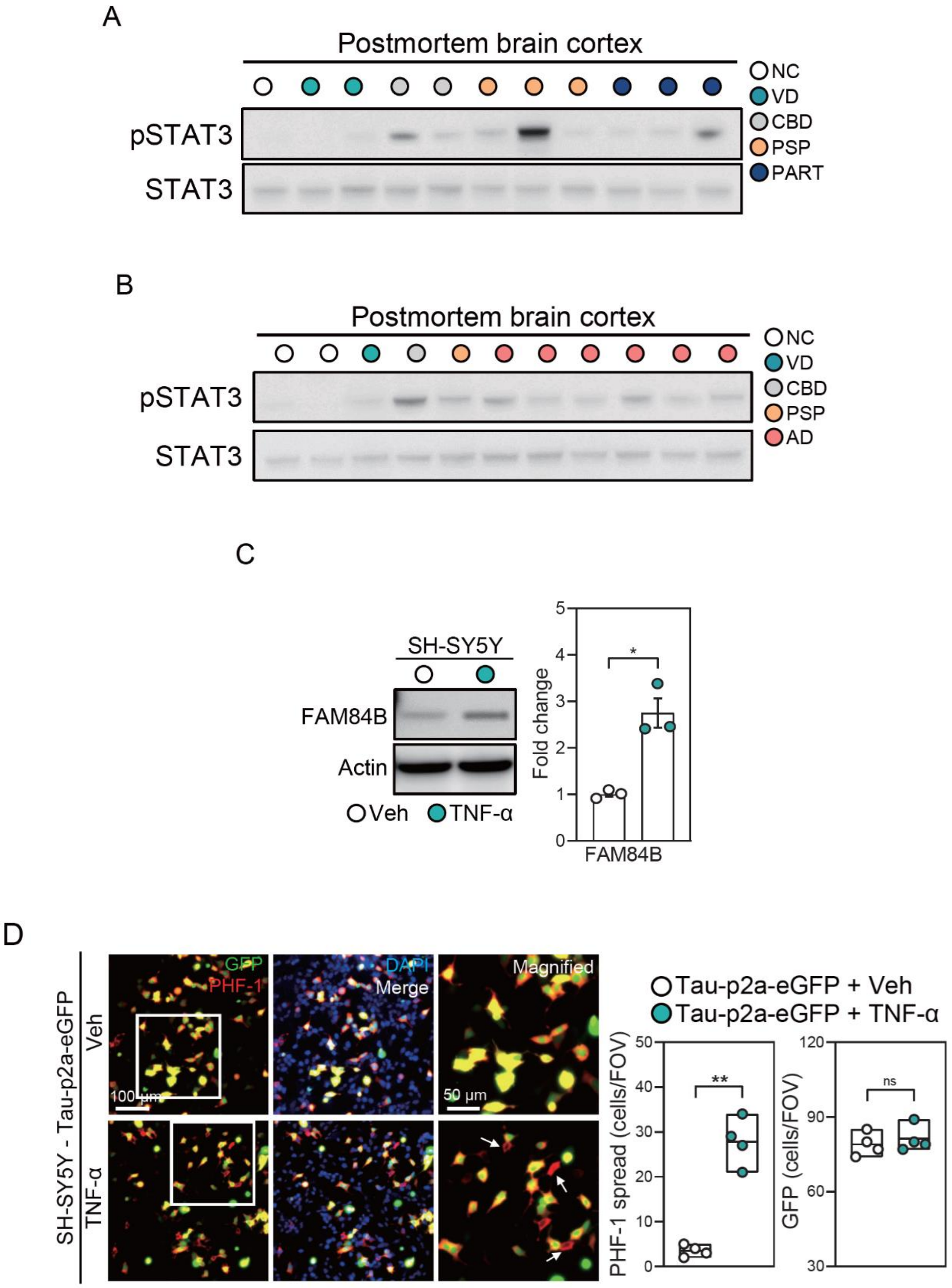

**Supplementary Figure 12**

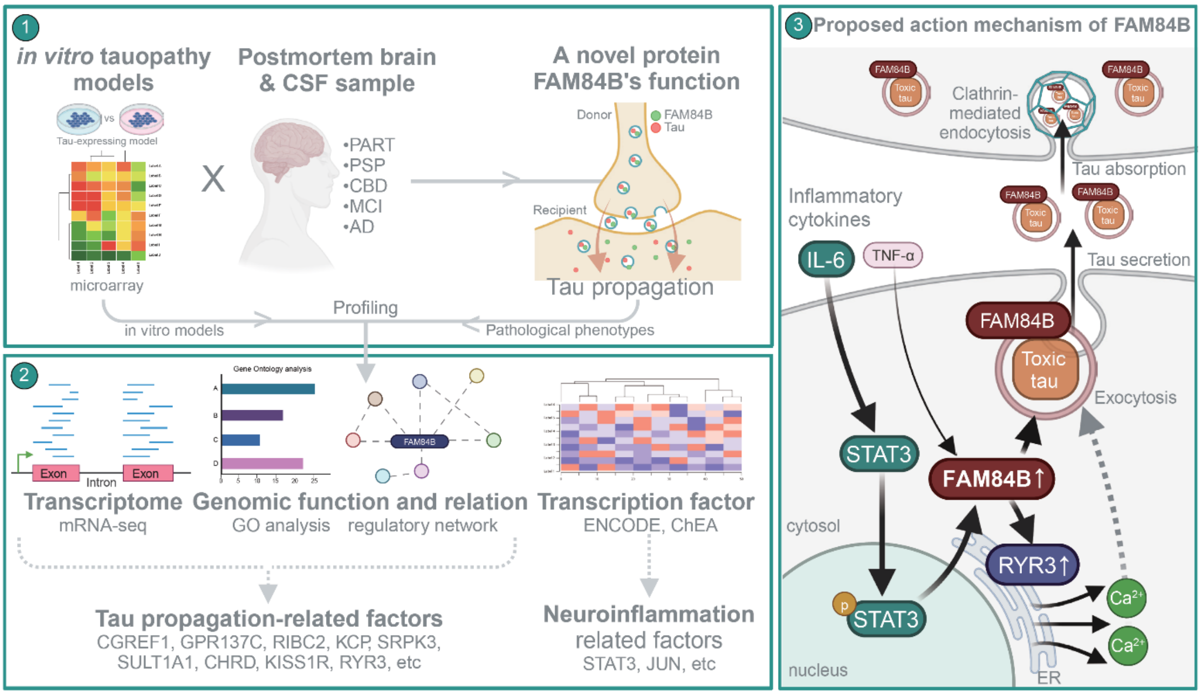

#### Supplementary Table 1

| primer name | Fw | Rev |
|---|---|---|
| pmCherry-C1-FAM84B | ATGGTACCATGGGCAACCAGGTGGAGAAAT | GAGAATTCTCAGTGTGCCACTGCCTCTC |
| pcDNA3.1-Tau-p2a-eGFP | CATGGATCCATGGCTGAGCCCCGCCAG | GACTCGAGCAAACCCTGCTTGGCCAGG |
| pAAV-FAM84B | TCGAATTCGATCCGGTACCGAGGAGAT | GAGGATCCTCAGTGTGCCACTGCCTCTC |
| pAAV-FAM84B-Myc | TCGAATTCGATCCGGTACCGAGGAGAT | GAGGATCCTCACAGATCCTCTTCTGAGATGAG |
| pAAV-Tau-Myc-p2a-eGFP | GGTGAATTCACCATGGCTGAGCCCCGCC | TGTCTAGACAGATCCTCTTCTGAGATGAG |

### Supplementary Table 2

| ENCODE project | CHEA prediction |
|---|---|
| ARID3A | AHR |
| ATF2 | AR |
| BACH1 | ARNT |
| BCL3 | ATF3 |
| BHLHE40 | BMI1 |
| BRCA1 | DACH1 |
| CBX3 | EED |
| CCNT2 | ELF5 |
| CEBPB | ESR1 |
| CHD1 | ETS2 |
| CHD2 | FOXP3 |
| CREB1 | GATA2 |
| CTBP2 | JARID2 |
| CTCF | JUN |
| E2F4 | MITF |
| E2F6 | PAX3 |
| EGR1 | PHC1 |
| ELF1 | PHF8 |
| ELK1 | PPARD |
| EP300 | RNF2 |
| ESR1 | RUNX2 |
| EZH2 | SMAD2 |
| FOS | SMAD3 |
| FOSL2 | SOX2 |
| FOXA1 | SOX9 |
| FOXA2 | STAT3 |
| FOXP2 | SUZ12 |
| GABPA | TFAP2C |
| GATA3 | TP53 |
| GTF2F1 | TP63 |
| H2AFZ | XRN2 |
| HCFC1 | ZNF217 |
| HDAC1 | ZNF281 |
| HDAC2 | |
| HDAC6 | |
| HMGN3 | |
| HNF4G | |
| IRF1 | |
| JUN | |
| JUND | |
| KDM4A | |
| KDM5A | |
| KDM5B | |
| MAFK | |
| MAX | |
| MAZ | |
| MXI1 | |
| MYC | |
| MYOG | |
| NFIC | |
| NFYA | |
| NR2F2 | |
| NR3C1 | |
| NRF1 | |
| PHF8 | |
| PML | |
| POLR2A | |
| PRDM1 | |
| RAD21 | |
| RBBP5 | |
| RCOR1 | |
| REST | |
| RFX5 | |
| RNF2 | |
| SAP30 | |
| SIN3A | |
| SMARCB1 | |
| SMC3 | |
| SP1 | |
| SP4 | |
| SRF | |
| STAT3 | |
| SUZ12 | |
| TAF1 | |
| TAF7 | |
| TBL1XR1 | |
| TBP | |
| TCF12 | |
| TCF7L2 | |
| TEAD4 | |
| TRIM28 | |
| UBTF | |
| USF1 | |
| USF2 | |
| YY1 | |
| ZBTB33 | |
| ZBTB7A | |
| ZEB1 | |
| ZKSCAN1 | |
| ZMIZ1 | |
| ZNF143 | |
| ZNF217 | |
| ZNF263 | |
| ZNF384 | |

**Supplementary Figure Legends**

**Supplementary Figure 1. Differential expression of FAM84B in tau-expressing cell models**

Gene expression changes in HEK293 cells expressing relatively high tau levels compared to low tau levels are shown in a heatmap and volcano plot (n=2 each). Genes with $p < 0.05$ and >10-fold expression change are marked with red dots in the volcano plot.

**Supplementary Figure 2. FAM84B expression changes in the brain tissue of patients with tauopathy**

**(A, B)** Immunoblots of FAM84B and actin in the cerebral cortex of HC and patients with VD, CBD, PSP, PART, and AD, supporting the data shown in Figure 1A.

HC, healthy controls; VD, vascular dementia; CBD, corticobasal degeneration; PSP, progressive supranuclear palsy; PART, primary age-related tauopathy; AD, Alzheimer's disease.

**Supplementary Figure 3. FAM84B overexpression decreases phosphorylated and total tau levels**

**(A)** Levels of PHF-1, total tau, FAM84B (Myc), and actin in HEK293 cells expressing varying amounts (0.2 µg, 0.4 µg, and 0.6 µg) of tau alone or tau with FAM84B (n=3 each).

**(B)** Levels of PHF-1, total tau, FAM84B (Myc), and actin in N2A cells expressing tau alone or tau with FAM84B-Myc, and in differentiated (Diff.) N2A and SH-SY5Y cells expressing control vector or FAM84B-Myc (n=3 each). The N2A and SH-SY5Y cells were treated with RA to induce differentiation.

Results are presented as mean ± SEM. Statistical analyses: Unpaired two-tailed *t*-test with Welch's correction; ns: not significant, $^{*}p < 0.05$, $^{**}p < 0.01$, $^{***}p < 0.001$.

PHF-1, phosphorylated tau (S396/S404); N2A, Neuro-2A; RA, retinoic acid.

**Supplementary Figure 4. FAM84B overexpression does not affect the transcription and degradation of tau proteins**

**(A)** MAPT mRNA expression in HEK293 cells expressing tau alone or tau with FAM84B-Myc (n=6 each).

**(B)** Levels of total tau, FAM84B (Myc), and actin in DOX-treated DIT cells expressing control vector or FAM84B-Myc (n=3 each).

**(C)** Levels of p62, LC3, and actin in HEK293 cells expressing tau alone or tau with FAM84B-Myc (n=3 each).

**(D)** Levels of HSP70, LAMP-1, LAMP-2, and actin in HEK293 cells expressing tau alone or tau with FAM84B-Myc (n=3 each).

Results are presented as mean ± SEM. Statistical analyses: unpaired two-tailed *t*-test with Welch's correction; ns: not significant; *$p < 0.05$.

Abbreviations: DIT, doxycycline-inducible tau-expressing; DOX, doxycycline.

**Supplementary Figure 5. FAM84B secretes oligomeric phosphorylated and total tau proteins**

**(A)** Representative immunoblots of mTau and FAM84B (Myc) in the CM from SH-SY5Y cells expressing tau alone or tau with FAM84B-Myc, supporting the data shown in Figure 1E (right panel; total tau quantification).

**(B)** Levels of PHF-1, total tau, FAM84B (Myc), and actin in HEK293 and SH-SY5Y cells expressing VN-P301 alone, VN-P301L with FAM84B-Myc, P301L-VC alone, and P301L-VC with FAM84B-Myc (n=3 each).

**(C)** Levels of phosphorylated tau (S396) in the CM from HEK293 and SH-SY5Y cells expressing VN-P301 alone, VN-P301L with FAM84B-Myc, P301L-VC alone, and P301L-VC with FAM84B-Myc (n=8 each).

**(D)** Levels of total tau and FAM84B (Myc) in the CM from HEK293 and SH-SY5Y cells expressing VN-P301 alone, VN-P301L with FAM84B-Myc, P301L-VC alone, and P301L-VC with FAM84B-Myc (n=3 each).

**(E)** Levels of PHF-1, total tau, FAM84B (Myc), and actin in SH-SY5Y cells expressing tau alone, tau with FAM84B-Myc, VN-P301 alone, VN-P301L with FAM84B-Myc, P301L-VC alone, and P301L-VC with FAM84B-Myc (n=3 each).

Results are presented as mean ± SEM. Statistical analyses: (B-E) Unpaired two-tailed *t*-test with Welch's correction; * $P < 0.05$, ** $P < 0.01$, *** $P < 0.001$.

Abbreviations: mTau, monomeric tau; CM, conditioned media; PHF-1, phosphorylated tau at residues S396/S404; VN-P301L, Venus N-terminus-tagged P301L mutant tau; P301L-VC, Venus C-terminus-tagged P301L mutant tau.

**Supplementary Figure 6. Phosphorylated and total tau proteins accumulate in FAM84B-depleted cells**

**(A)** Changes in the levels of FAM84B and actin in FAM84B KO HEK293 cells compared to controls (n=3 each).

**(B)** Levels of total tau and FAM84B in the CM from normal or FAM84B KO HEK293 cells expressing wild-type tau, VN-P301L, and P301L-VC (n=3 each).

**(C)** Levels of phosphorylated tau (S396) in the CM from normal or FAM84B KO HEK293 cells expressing wild-type tau, VN-P301L, and P301L-VC (n=8 each).

**(D)** Levels of PHF-1, total tau, FAM84B, and actin in normal or FAM84B KO HEK293 cells expressing wild-type tau, VN-P301L, and P301L-VC (n=3 each).

Results are presented as mean ± SEM. Statistical analyses: Unpaired two-tailed *t*-test with Welch's correction; *$p < 0.05$, **$p < 0.01$, ***$p < 0.001$.

Abbreviations: KO, knockout; CM, conditioned media; VN-P301L, Venus N-terminus-tagged P301L mutant tau; P301L-VC, Venus C-terminus-tagged P301L mutant tau; PHF-1, phosphorylated tau at S396/S404 residue.

**Supplementary Figure 7. FAM84B colocalizes with phosphorylated and total tau proteins**

**(A)** Colocalization (arrow) of tau and FAM84B in HEK293 cells expressing GFP-tau with mCherry-FAM84B.

**(B)** Colocalization (arrow) of tau and FAM84B in differentiated (Diff.) SH-SY5Y cells expressing GFP-tau with mCherry-FAM84B.

**(C)** Colocalization (arrow) of tau and FAM84B in SH-SY5Y cells expressing tau with FAM84B-Myc.

**(D)** Colocalization (arrow) of tau and FAM84B in HEK293 cells expressing tau with FAM84B-Myc.

**(E)** Colocalization (arrow) of PHF-1 and FAM84B in SH-SY5Y cells expressing tau with FAM84B-Myc.

**(F)** Colocalization (arrow) of PHF-1 and FAM84B in HEK293 cells expressing tau with FAM84B-Myc.

**(G)** Histogram showing colocalization data from (A-F) using Pearson's correlation analysis. Results are presented as mean ± SEM (n=4 each).

Abbreviations: PHF-1, phosphorylated tau at S396/S404 residue.

**Supplementary Figure 8. FAM84B encapsulates tau proteins via cell membrane-associated vesicle formation**

**(A)** Immunoblots depicting the interaction between tau and FAM84B (Myc) in HEK293 cells expressing tau alone or tau with FAM84B-Myc (under detergent and non-reducing conditions).

**(B)** Immunoblots illustrating the interaction between tau and FAM84B (Myc) in conditioned media from HEK293 cells expressing tau alone or tau with FAM84B-Myc (under non-detergent and non-reducing conditions).

**(C)** Immunoblots demonstrating the interaction between tau and FAM84B (Myc) in conditioned media from HEK293 cells expressing tau alone or tau with FAM84B-Myc (under non-detergent and reducing conditions).

**(D)** Immunoblots presenting the interaction between tau and FAM84B (Myc) in conditioned media from HEK293 cells expressing tau alone or tau with FAM84B-Myc (under detergent and non-reducing conditions).

Immunoblots shown in (A–D) are representative of three independent experiments.

**Supplementary Figure 9. FAM84B is taken up together with phosphorylated and total tau proteins**

**(A)** Levels of PHF-1, total tau, FAM84B (Myc), and actin in HEK293 and SH-SY5Y cells exposed to the CM from normal cells, cells expressing tau alone, or cells expressing tau with FAM84B-Myc (n=3 each).

**(B)** Uptake and colocalization of total tau and FAM84B in HEK293 cells exposed to the CM from normal cells, cells expressing tau alone, or cells expressing tau with FAM84B-Myc (n=4 each).

**(C)** Uptake and colocalization of total tau and FAM84B in SH-SY5Y cells exposed to the CM from normal cells, cells expressing tau alone, or cells expressing tau with FAM84B-Myc (n=4 each).

Results are presented as mean ± SEM. Co-localization analysis: (B, C) Pearson's correlation coefficient. Statistical analyses: Unpaired two-tailed *t*-test with Welch's correction; ns: not significant, $*p < 0.05$, $**p < 0.01$, $***p < 0.001$.

Abbreviations: PHF-1, phosphorylated tau at residues S396/S404; CM, conditioned media.

**Supplementary Figure 10. FAM84B induces tau propagation via the increase of RYR3-mediated cytosolic calcium**

**(A)** Colocalization (arrow) of FAM84B and calcium in HEK293 cells expressing tau with mCherry control (C1), and in cells expressing tau with mCherry-FAM84B treated with either DMSO or dantrolene (n=4 each).

**(B)** PHF-1 spread from Tau-p2a-eGFP expressing HEK293 cells to neighboring cells in response to treatment with either DMSO or dantrolene (n=4 each). Arrows indicate recipient cells (red) to which tau has been transmitted. Solid boxes in the images indicate magnified areas.

Results are presented as mean ± SEM. Co-localization analysis: (A) Pearson's correlation coefficient. Statistical analyses: unpaired two-tailed *t*-test with Welch's correction; ns: not significant; ***$p < 0.001$.

Abbreviations: PHF-1, phosphorylated tau at S396/S404 residue.

**Supplementary Figure 11. The pro-inflammatory cytokine TNF-α also increases FAM84B expression and induces tau propagation**

**(A, B)** Immunoblots of phosphorylated STAT3 (Y705) and STAT3 in the cerebral cortex of HC and patients with VD, CBD, PSP, PART, and AD, supporting the data shown in Figure 5C.

**(C)** Levels of FAM84B and actin in SH-SY5Y cells treated with either control vehicle or TNF-α (n=3 each).

**(D)** PHF-1 spread from Tau-p2a-eGFP expressing SH-SY5Y cells to neighboring cells in response to treatment with either control vector or TNF-α (n=4 each). Arrows indicate recipient cells (red) to which tau has been transmitted. Solid boxes in the images indicate magnified areas.

Results are presented as mean ± SEM. Statistical analyses: (C, D) Unpaired two-tailed *t*-test with Welch's correction; ns: not significant, * $P < 0.05$, ** $P < 0.01$.

Abbreviations: HC, healthy controls; VD, vascular dementia; CBD, corticobasal degeneration; PSP, progressive supranuclear palsy; PART, primary age-related tauopathy; AD, Alzheimer's disease; PHF-1, phosphorylated tau at residues S396/S404

**Supplementary Figure 12. FAM84B plays a key role in tau propagation in tauopathy**

**(1)** Identification of the role of FAM84B in tau propagation using various tau-expressing cell models (including those analyzed by microarray), postmortem brain tissue (PART, PSP, CBD, MCI, and AD), and CSF from patients with tauopathies.

**(2)** Investigation of the mechanism in tau propagation mediated by FAM84B through transcriptome analysis (mRNA-seq), functional gene identification (GO analysis), and transcription cofactor prediction (ChIP-seq data from ENCODE, ChEA).

**(3)** Proposed mechanism of action of FAM84B: Upon cellular stress, such as that associated with the release of inflammatory cytokines (IL-6 and TNF-α), activated STAT3 directly upregulates FAM84B levels which in turn induces the expression of RYR3, leading to altered cytosolic calcium concentration. This process facilitates the encapsulation of toxic tau by the membrane protein FAM84B and its subsequent secretion. The released exosomal tau is taken up by neighboring cells via clathrin-mediated endocytosis, thereby propagating pathological tau.

Illustration was created with BioRender.com (Agreement number: OF2789J8GK)

Abbreviations: PART, primary age-related tauopathy; PSP, progressive supranuclear palsy; CBD, corticobasal degeneration; MCI, mild cognitive impairment; AD, Alzheimer's disease; CSF, cerebrospinal fluid; GO, gene ontology.

**Supplementary Table Legends**

The supplementary table files are available online.

**Supplementary Table 1. List of primers used for subcloning**

The sequences of the primer sets used for subcloning are listed. Primers containing restriction enzyme sites for insertion into vectors were included.

**Supplementary Table 2. The data from bioinformatics analyses**

Genes selected from the mRNA-seq databases were included. The cellular compartment terms of Gene Ontology (GO) analysis were included in the order of a significant number of genes. The predicted transcription factors and chromatin regulators were included.
